# Molecular basis of collagen triple helix recognition by VWF A-like domain 2 of collagen VII: Implications for interlaced anchoring fibril formation

**DOI:** 10.64898/2026.03.16.711976

**Authors:** Moe Hashimoto, Hiroya Oki, Kazuki Kawahara, Kazunori K. Fujii, Takaki Koide

**Affiliations:** Department of Chemistry and Biochemistry, School of Advanced Science and Engineering, Waseda University, Shinjuku, Tokyo 169-8555, Japan; Department of Infection Metagenomics, Genome Information Research Center, Research Institute for Microbial Diseases, The University of Osaka, Suita, Osaka 565-0871, Japan; Graduate School of Pharmaceutical Sciences, The University of Osaka, Suita, Osaka 565-0871, Japan

**Author notes:** Corresponding author: Takaki Koide, Ph.D. Professor, Department of Chemistry and Biochemistry, School of Advanced Science and Engineering, Waseda University, Shinjuku, Tokyo 169-8555, Japan; Kazuki Kawahara, Ph.D. Assistant Professor, Graduate School of Pharmaceutical Sciences, The University of Osaka, Suita, Osaka 565-0871, Japan.

## Abstract

Anchoring fibrils formed by collagen VII play a critical role in stabilizing the dermal–epidermal junction. The N-terminal non-collagenous (NC1) domain of collagen VII binds firmly to basement membrane components including collagen IV and has also been reported to interact with mesenchymal fibrillar collagens *via* its von Willebrand factor A-like domain 2 (A2 domain). To elucidate how collagen VII recognizes fibrillar collagen, we performed yeast two-hybrid screening using a triple-helical random peptide library, which resulted in the identification of a Met-Gly-Φ (Φ; aromatic amino acid residue) motif. Biochemical analysis with synthetic triple-helical peptides revealed a binding preference of Trp > Phe as the Φ residue by the A2 domain despite Trp being absent in native collagens. The crystal structure of the A2 domain in complex with the Nle (Met surrogate)-Gly-Trp-containing peptide revealed a unique mechanism by which two distinct hydrophobic pockets of the A2 domain accommodate the Nle and Trp residues corresponding to the Met-Gly-Φ motif, engaging all three chains of the triple helix. Subsequent molecular dynamics simulations demonstrated that the A2 domain recognizes the corresponding native Met-Gly-Phe motif in a similar manner, but with lower affinity, implying a transient interaction with mesenchymal collagens. The findings obtained in this work suggest models in which transient A2-triple helix interaction promotes the recruitment of collagen I and III fibrils into the arc-shaped structure of anchoring fibrils. This also provides a foundation for linking structural understanding to skin fragility diseases caused by collagen VII dysfunction.

## Introduction

The dermal–epidermal junction (DEJ) secures the epidermis to the dermis, thereby maintaining skin structural stability (Figure 1A). The DEJ consists of four major components: hemidesmosomes on the basal surface of keratinocytes, anchoring filaments that connect hemidesmosomes to the basement membrane, the basement membrane itself, and anchoring fibrils that link the basement membrane to the dermal extracellular matrix (ECM) (1).

**Figure 1.**
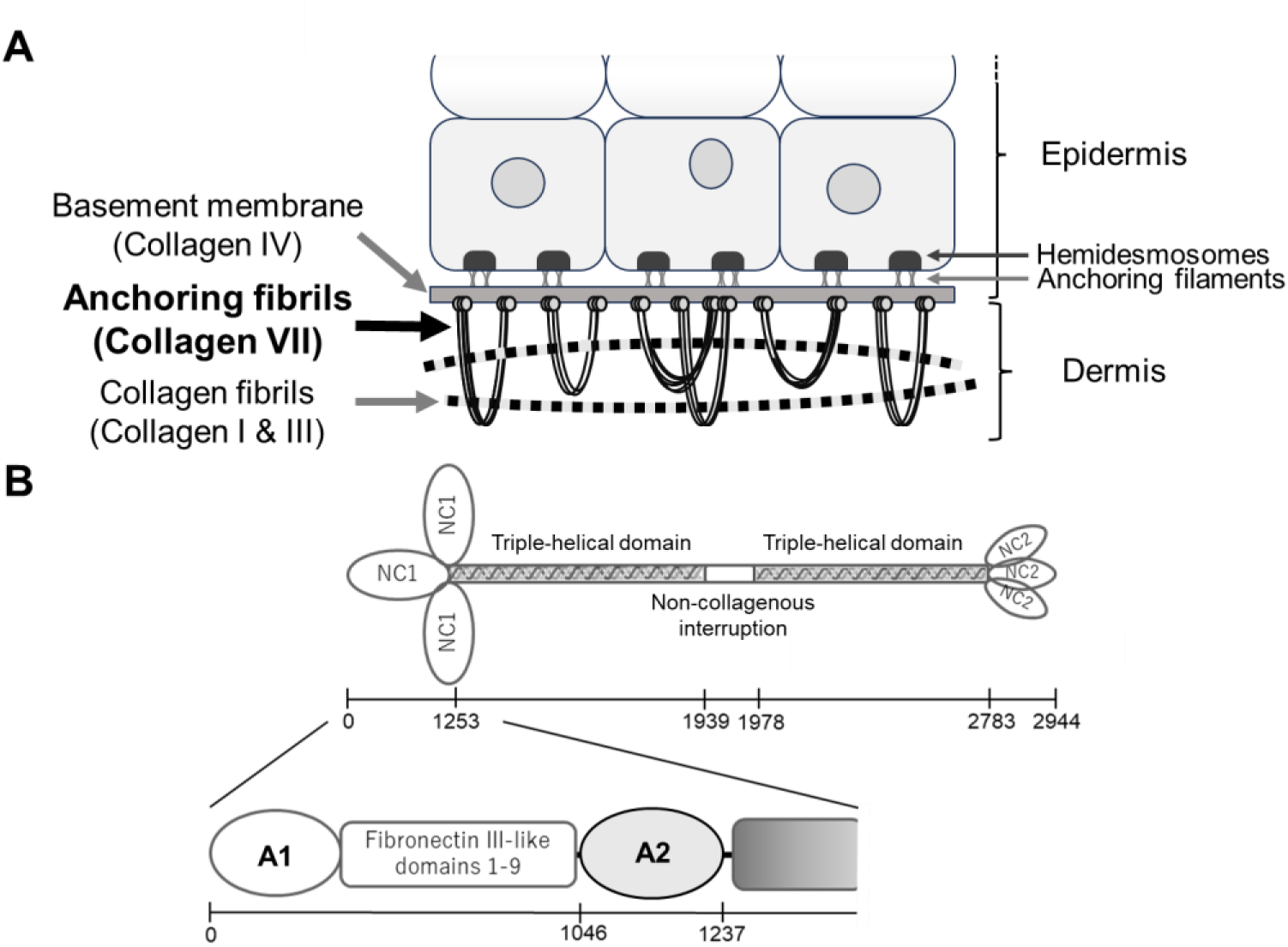
Structure of the DEJ and schematic representation of collagen VII. *A,* Collagen VII forms anchoring fibrils and interacts with collagens I, III, and IV at the DEJ. *B,* Schematic representation of a triple-helical collagen VII molecule. The triple-helical domain is flanked by NC1 and NC2 domains and contains a 39-residue non-collagenous interruption. NC1 domain contains von Willebrand factor A-like domain 1 (A1domain) and von Willebrand factor A-like domain 2 (A2 domain).

Anchoring fibrils exhibit a symmetric arc-like structure with both ends embedded in the basement membrane. Collagen VII constitutes anchoring fibrils by forming a unique supramolecular arc-like architecture (2). Collagen I and III fibrils have been observed traversing through these arcs (3–5). Mutations in *COL7A1* cause deficiency or structural abnormalities of anchoring fibrils, leading to dystrophic epidermolysis bullosa (DEB). DEB is characterized by skin fragility, blistering, and erosion (6–9). A clinically related acquired autoimmune disorder, epidermolysis bullosa acquisita (EBA), is caused by IgG autoantibodies directed against collagen VII (10,11).

Collagens are prominent ECM proteins characterized by a triple-helical structure composed of Gly-Xaa-Yaa repeating sequences, where Xaa is proline and Yaa is 4(*R*)-hydroxyproline (Hyp, O). Hyp is generated by post-translational modification of proline and contributes to the conformational stability of the triple helix (12,13). Collagen VII is a homotrimer of three α chains, consisting of a central triple-helical domain flanked by the N-terminus non-collagenous domain (NC1) and the C-terminus non-collagenous domain (NC2). The triple-helical region contains a 39-residue non-collagenous interruption (Figure 1B) (14,15). Triple-helical collagen VII molecules assemble into antiparallel dimers through proteolytic cleavage of a specific region within the NC2 domain (16,17). Over 10 dimeric molecules subsequently laterally aggregate to form anchoring fibrils with NC1 domains at both ends (18,19).

In the mature DEJ, the NC1 domain is tightly connected to basement membrane components such as collagen IV and laminin 332 (20–23). Collagen IV contains not only a triple-helical region but also a globular NC1 domain. Compared with its triple-helical region, the globular NC1 domain of collagen IV was reported to mediate stronger interactions with collagen VII NC1 domain (20). Triple-helical regions of collagen I and III, which traverse through collagen VII arcs, were also reported to bind to collagen VII NC1 domain (21,24–26). The NC1 domain of collagen VII is organized into 11 subdomains. Two of these subdomains, the von Willebrand factor A-like domain 1 and von Willebrand factor A-like domain 2, are homologous to the von Willebrand factor A3 domain (VWF A3) and flank fibronectin type III-like repeats (14). These two domains are hereafter referred to as A1 and A2 domains respectively. The A2 domain has been shown to mediate interactions between collagen VII and other ECM proteins, including triple helices of collagen I and III (24,26,27). This domain also represents the major autoantigen in EBA (28–30). Although these interactions explain how collagen VII is anchored to basement membrane, they do not clarify how collagen I and III fibrils become engaged by the arched rings of the anchoring fibrils. In particular, the molecular basis of how A2 recognizes collagen triple helix remains unclear.

Our research group has developed a yeast two-hybrid (Y2H) screening system employing a triple-helical random peptide library that enables the identification of triple-helical sequences recognized by collagen-binding proteins (31,32). Using this system, we identified collagen triple-helical sequences recognized by the A2 domain and elucidated the molecular basis of its interaction with the triple-helical peptide. Furthermore, defining the human collagen sequences bound to the A2 domain provided insights into the forming process of collagen VII anchoring fibrils in which mesenchymal collagen fibrils are threaded through.

## Results

### Selection of collagen VII A2-binding peptides from a random peptide library

To identify triple-helical peptides that interact with collagen VII, we employed Y2H screening of a random peptide library. In this setup, A1 domain or A2 domain of the human collagen VII was expressed as a fusion protein with the GAL4 DNA-binding domain (GAL4-BD) and served as bait, while the peptide library was fused to the GAL4 activation domain (GAL4-AD). The library was designed by randomizing six residues within the collagen-like sequence Yaa1-Gly2-Xaa3-Yaa4-Gly5-Xaa6-Yaa7-Gly8-Xaa9 as described in a previous study (31).

The random peptide library contained 7.34 × 10⁵ clones. The screening for A1-binding sequences yielded no positive colonies, whereas screening for the A2-binding peptides yielded 12 independent clones (Table 1). All identified sequences contained Met at the Yaa1 position. At the Xaa3 position, aromatic residues such as Phe, Trp, and Tyr were highly enriched, yielding a common Met-Gly-Φ (Φ; aromatic amino acid residue) motif. Clone 5 was the sole exception in which Pro occupied the Xaa3 position. However, this clone contained Met at the Yaa4 position and Trp at the Xaa6 position, thereby maintaining the Met-Gly-Φ signature. None of the 12 identified sequences matched any known human collagen α-chain sequences.

**Table 1.** Collagen VII A2-binding sequences identified from the peptide library.

| Name | Sequence | Frequency |
| --- | --- | --- |
|  | Yaa1-Gly2-Xaa3-Yaa4-Gly5-Xaa6-Yaa7-Gly8-Xaa9 |  |
| Clone 1 | Met-Gly-Phe-Gln-Gly-Leu-Arg-Gly-Asp | 13/33 |
| Clone 2 | Met-Gly-Trp-Trp-Gly-Pro-Ser-Gly-Thr | 5/33 |
| Clone 3 | Met-Gly-Tyr-Trp-Gly-Arg-Lys-Gly-Ser | 4/33 |
| Clone 4 | Met-Gly-Phe-Met-Gly-Ala-Met-Gly-Ala | 3/33 |
| Clone 5 | Met-Gly-Pro-Met-Gly-Trp-Met-Gly-Leu | 1/33 |
| Clone 6 | Met-Gly-Phe-Met-Gly-Asn-His-Gly-Ser | 1/33 |
| Clone 7 | Met-Gly-Trp-Met-Gly-Ala-Lys-Gly-Arg | 1/33 |
| Clone 8 | Met-Gly-Trp-Trp-Gly-Ala-Thr-Gly-Thr | 1/33 |
| Clone 9 | Met-Gly-Phe-Met-Gly-Leu-Arg-Gly-Phe | 1/33 |
| Clone 10 | Met-Gly-Phe-Trp-Gly-Arg-Lys-Gly-Ser | 1/33 |
| Clone 11 | Met-Gly-Phe-Gln-Gly-Ser-Arg-Gly-Met | 1/33 |
| Clone 12 | Met-Gly-Phe-Trp-Gly-Pro-Ile-Gly-Gly | 1/33 |

To evaluate interaction with the A2 domain, three representative sequences were selected. Clone 1 was the most abundant sequence in the selection. Clone 4 contained three Met residues at Yaa positions. Clone 7 was predicted to form the most thermally stable triple helix among the sequences containing Trp at Xaa3 (Table S2) (33). Peptides containing these sequences were tested for interaction with A2 in the Y2H system under the same conditions as the initial screening (Figure S1). We also examined their binding to heat-shock protein 47 (HSP47). HSP47 is a molecular chaperone serving as an indicator of the triple-helical conformation in the Y2H system (31,34,35). As controls, a peptide containing a pigment epithelium-derived factor (PEDF)-binding sequence (Lys-Gly-His-Arg-Gly-Phe-Ser-Gly-Leu, Seq *a*) was used (36). All three peptides corresponding to Clones 1, 4, and 7, identified in the Y2H screening, interacted with the A2 domain but not with PEDF, and each bound to HSP47. The observed HSP47 binding confirms that these peptides adopt a triple-helical conformation at the assay temperature of 25°C.

### Evaluation of relative collagen VII A2 domain-binding affinity using synthetic triple-helical peptides

Peptides Pep 1, Pep 2, and Pep 3 corresponding to the sequences derived from Clone 1, Clone 4, and Clone 7, respectively, were chemically synthesized for further biochemical examinations. Each peptide was designed as an open-chain homotrimeric triple helix with the sequence H-(Pro-Hyp-Gly)₄-Pro-[guest sequence]-Hyp-Gly-(Pro-Hyp-Gly)₄-Pro-Tyr-NH₂.

In search for natural triple-helical sequences potentially recognized by the A2 domain, we searched for human collagen sequences with the Met-Gly-Φ signature, which were obtained in the screening (Table 2). Ten sequences were found to match it, seven containing Phe and three containing Tyr as the aromatic residue. Notably, none of the natural sequences contained Trp, which was frequently present in the screening-derived clones. Among these sequences, we selected a collagen III region containing the Met-Gly-Φ motif for additional peptide synthesis (Pep 4). This peptide was selected due to the homotrimeric structure of collagen III, which facilitates its chemical preparation and comparison with other peptides (Table 3).

**Table 2.**
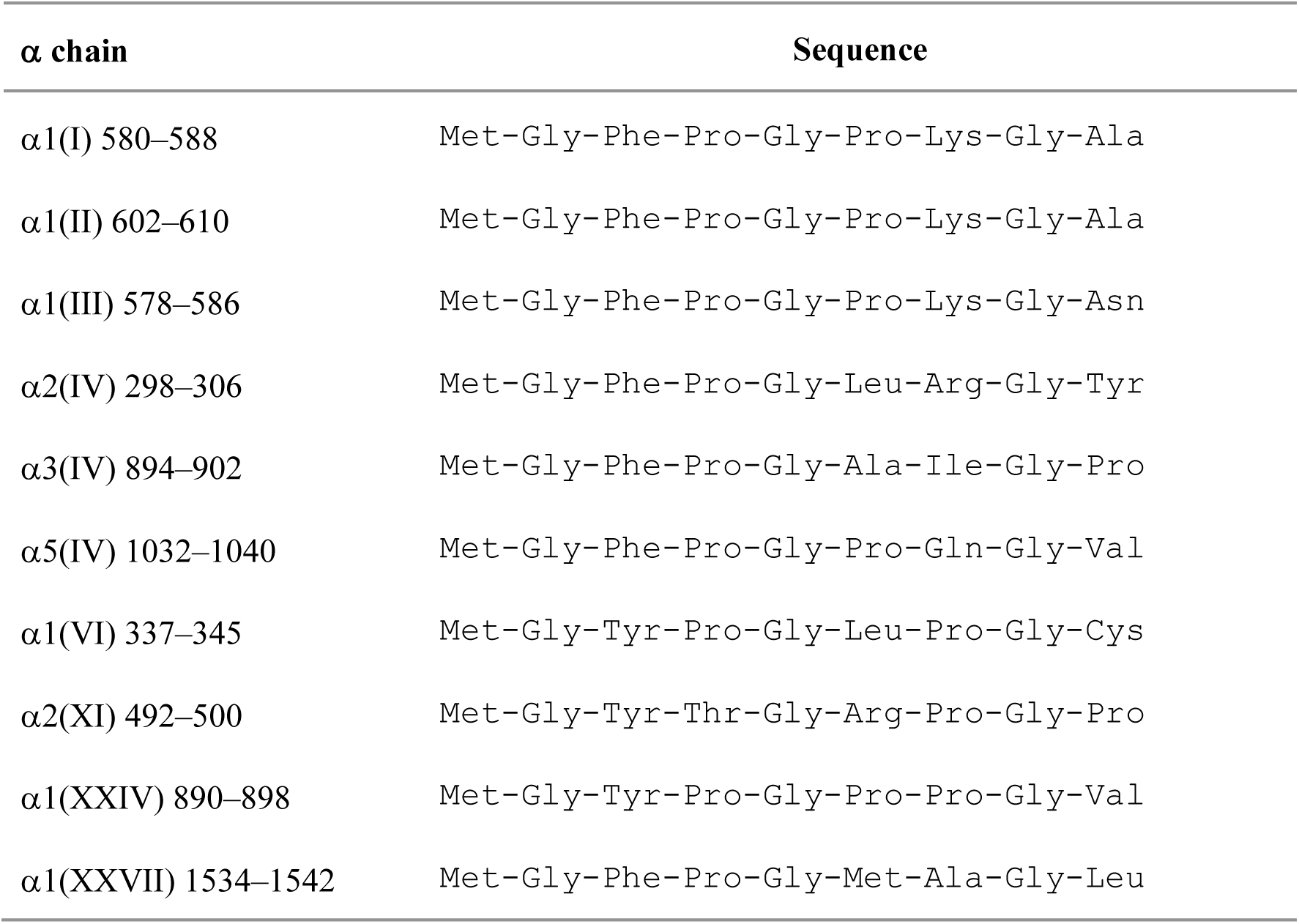
Human collagen sequences containing Met-Gly-Φ signature.

**Table 3.** Sequences of synthesized triple-helical peptides of the format H-(Pro-Hyp-Gly)4-Pro-[guest sequence]-Hyp-Gly-(Pro-Hyp-Gly)4-Pro-Tyr-NH2.

| Name | Guest sequence | Origin |
| --- | --- | --- |
| Pep 1 | Met-Gly-Phe-Gln-Gly-Leu-Arg-Gly-Asp | Clone 1 |
| Pep 2 | Met-Gly-Phe-Met-Gly-Ala-Met-Gly-Ala | Clone 4 |
| Pep 3 | Met-Gly-Trp-Met-Gly-Ala-Lys-Gly-Arg | Clone 7 |
| Pep 4 | Met-Gly-Phe-Hyp-Gly-Pro-Lys-Gly-Asn | human $\alpha 1(\text{III})$ 578–586 |

The relative binding affinities of the peptides listed in Table 3 for collagen VII A2 domain were investigated. An enzyme-linked immunosorbent assay (ELISA) was conducted using peptide polymers coated onto wells and recombinantly expressed glutathione *S*-transferase (GST)-tagged collagen VII A2 domain. Triple-helical peptide polymers were prepared from peptides bearing three Cys residues at each terminus *via* disulfide bond-formation as described previously (37). C3-Pep 4 contains the same guest sequence as Pep 4, whereas C3, which includes the Gly-Pro-Hyp-Gly-Pro-Arg sequence, served as a control (Table S1). These peptides were used for preparing corresponding polymers. Binding of the GST-tagged A2 domain increased with the proportion of C3-Pep 4 in the peptide polymer (Figure S6). GST-collagen VII A2 binding to C3-Pep 4 polymer was examined in the presence of soluble peptides listed in Table 3 as competitors (Figure 2A). Pep 0, H-(Pro-Hyp-Gly)6-Pro-Arg-Gly-(Pro-Hyp-Gly)5-Pro-Tyr-NH2, was also used as a control (Table S1). Among the four peptides tested, Pep 1 containing MGFQGLRGD and Pep 4 containing MGFOGPKGN showed comparable inhibiting concentration for the A2 domain. Pep 2 containing MGFMGAMGA inhibited the A2 binding at concentrations approximately five-fold lower than these peptides. Notably, Pep 3 required an approximately 100-fold lower concentration than Pep 1 and Pep 4. This result suggested that Trp, as the Φ residue in the Met-Gly-Φ motif, confers higher-affinity binding to the A2 domain than Phe.

**Figure 2.**
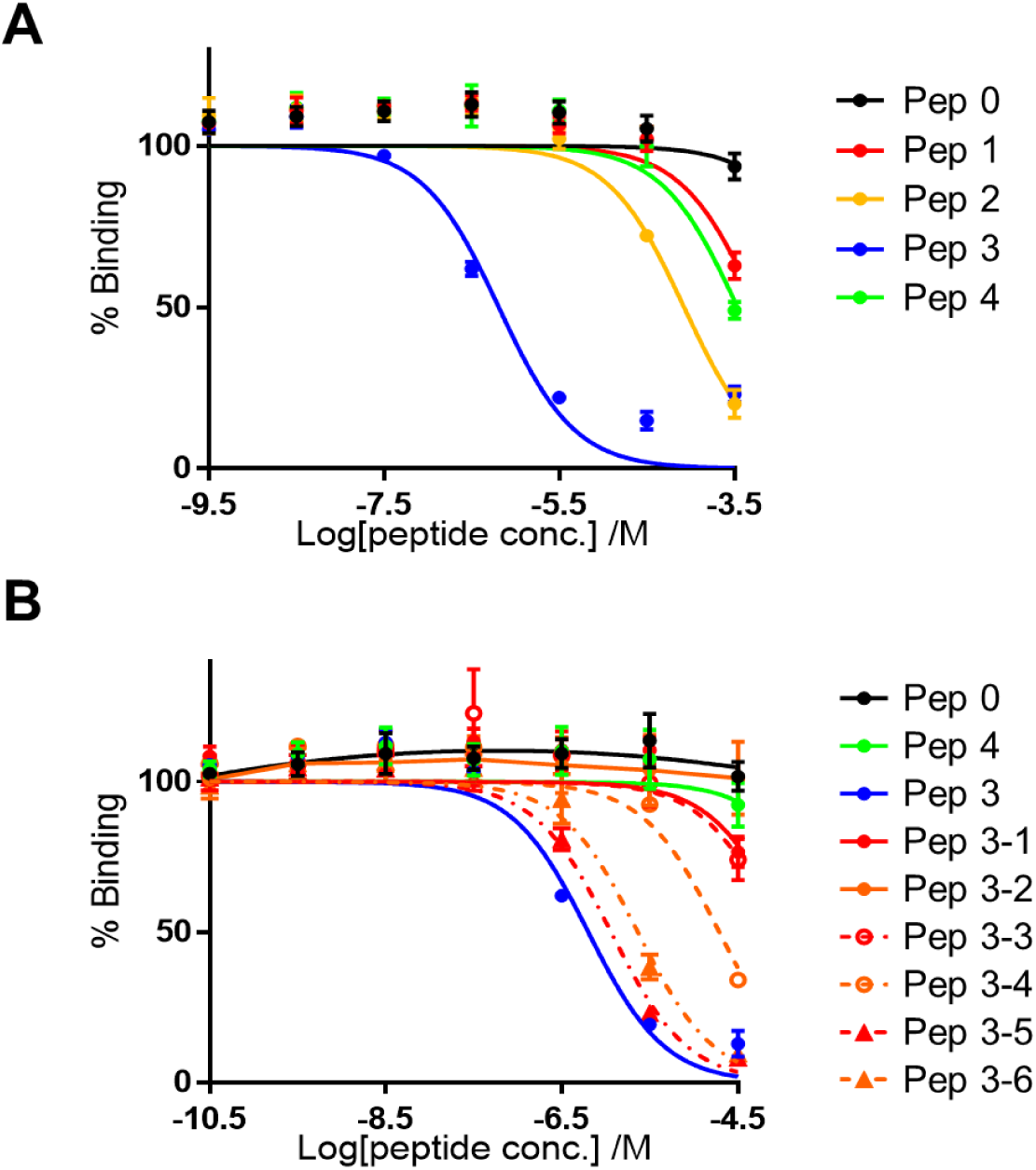
Evaluation of the relative binding affinity of triple-helical peptides for collagen VII A2. *A*, Binding of 10 nM GST-collagen VII A2 to wells coated with 100% C3-Pep 4 polymer (0.5 μg/well) was inhibited by the peptides listed in Table 3. The ELISA was conducted at 25°C (n = 3, mean ± S.D.). *B*, Binding of 10 nM GST-collagen VII A2 to the wells coated with 100% C3-Pep 4 polymer (0.5 μg/well) was inhibited by Pep 3 and its amino acid substitution variants. The experiment was conducted at 25°C (n = 3, mean ± S.D.).

To identify the amino acid residues in Pep 3 that are critical for recognition by collagen VII A2 domain, a structure–activity relationship analysis was conducted (Figure 2B). Each residue of Pep 3 was substituted individually with Pro at the Xaa position or Hyp at the Yaa position (Table S1), and the inhibitory activity of the synthetic peptides were evaluated using the similar competitive ELISA. Because Gly-Pro-Hyp triplets constitute the common repeat unit of a triple helix, Xaa-Pro and Yaa-Hyp scanning allow identification of residues critical for binding, without disrupting the triple-helical structure. The IC50 values for each peptide are summarized in Table 4. The greatest increase in IC50 was observed when Trp at Xaa3 was replaced by Pro (Pep 3-2), corresponding to more than a 10,000-fold increase relative to the parent peptide Pep 3. Substitution of Met residues at Yaa1 or Yaa4 with Hyp (Pep 3-1 and Pep 3-3, respectively) led to more than a 100-fold increase in IC50, while replacement of Ala at Xaa6 with Pro (Pep 3-4) caused approximately a 30-fold increase. Substitution of Lys at Yaa7 (Pep 3-5) or Arg at Xaa9 (Pep 3-6) resulted in smaller increases in IC50 compared with substitutions at other positions. These results indicate that MGWMGA sequence of Pep 3 is particularly important for A2 recognition. This result is consistent with the result shown in Figure 2A, highlighting the particular contribution of the Trp residue to the high-affinity binding of the A2 domain.

**Table 4.** IC50 values of Pep 3 and its Pro- and Hyp-scanning variants.

| Peptide | Guest sequence | IC<sub>50</sub> (μM) <sup>a</sup> |
| --- | --- | --- |
|  | Yaa1-Gly2-Xaa3-Yaa4-Gly5-Xaa6-Yaa7-Gly8-Xaa9 |  |
| Pep 0 | Hyp-Gly-Pro-Hyp-Gly-Pro-Arg-Gly-Pro | $>3.3 \times 10^4$ |
| Pep 4 | Met-Gly-Phe-Hyp-Gly-Pro-Lys-Gly-Asn | $(4.5 \pm 4.2) \times 10^2$ |
| Pep 3 | Met-Gly-Trp-Met-Gly-Ala-Lys-Gly-Arg | $0.63 \pm 0.12$ |
| Pep 3-1 | **Hyp**-Gly-Trp-Met-Gly-Ala-Lys-Gly-Arg | $(1.9 \pm 0.5) \times 10^2$ |
| Pep 3-2 | Met-Gly-**Pro**-Met-Gly-Ala-Lys-Gly-Arg | $>3.3 \times 10^4$ |
| Pep 3-3 | Met-Gly-Trp-**Hyp**-Gly-Ala-Lys-Gly-Arg | $94 \pm 35$ |
| Pep 3-4 | Met-Gly-Trp-Met-Gly-**Pro**-Lys-Gly-Arg | $20 \pm 4$ |
| Pep 3-5 | Met-Gly-Trp-Met-Gly-Ala-**Hyp**-Gly-Arg | $1.1 \pm 0.1$ |
| Pep 3-6 | Met-Gly-Trp-Met-Gly-Ala-Lys-Gly-**Pro** | $2.3 \pm 0.3$ |

<sup>a</sup> IC<sub>50</sub> values are means ± S.E.M.

### Structural basis of collagen VII A2 recognition of triple-helical peptide containing the Met-Gly-Φ motif

The screening result and the structure–activity relationship analyses in the previous section have highlighted the importance of the Met-Gly-Φ motif in the recognition by the human collagen VII A2 domain. To elucidate the molecular recognition mechanism, we performed an X-ray crystallographic analysis of the A2 domain in complex with the triple-helical peptide containing the sequence Ac-(Gly-Pro-Hyp)2-Gly-Pro-Nle-Gly-Trp-Nle-Gly-Ala-Hyp-(Gly-Pro-Hyp)2-NH2 (Short-XGWXGA) at 1.47 Å resolution (Tables S1 and S3, Figure S7). This peptide contains the XGWXGA motif, which forms the minimal binding motif of Pep 3-X (Table S1), the norleucine (Nle, X)-substituted variant of Pep 3 that exhibits the highest affinity for A2 among those tested (Figure S8). Because Met residues are susceptible to oxidation, Nle residues which have the same side-chain length but are resistant to oxidation were used. The peptide Short-XGWXGA comprises 21 amino acid residues. An attempt to co-crystallize A2 domain with a longer peptide containing additional Gly-Pro-Hyp triplets at both termini was unsuccessful.

As shown in Figure 3A, Short-XGWXGA binds along the β3 strand of the A2 domain. The A2 structure in the complex was nearly identical to that of the mouse apo form (PDB: 6S4C) (38), with a root-mean-square deviation (RMSD) of 0.506 Å, indicating no significant conformational change upon ligand binding (Figure S9). Two hydrophobic pockets flanking the β3 strand of the A2 domain respectively accommodated the side chains of Nle9 and Trp11 residues in the Short-XGWXGA (Figure 3B). The first pocket (Pocket 1) accommodated Trp11 from the trailing chain, together with Nle9 and Nle12 from the middle chain. The second pocket (Pocket 2) was occupied by Trp11 of the middle chain of the triple helix, along with Nle9 from the leading chain, and Pro8 from the middle chain. Trp11 of the middle chain engaged in CH–π interactions with Pro58 and Pro61 of the A2 domain (Figure 3C). While such peptide recognition by the A2 domain would be primarily driven by hydrophobic interactions, several hydrogen bonds also contributed to the binding (Figure 3D). Notably, Trp11 of the trailing chain formed a hydrogen bond *via* the Nε1 atom of its indole ring with Ser57. Trp11 was identified in the Pro-and Hyp-scanning analysis as a position whose substitution markedly increased the IC50 (Figure 2, Table 4). In addition, Gly13 of the middle chain formed a hydrogen bond with Arg74.

**Figure 3.**
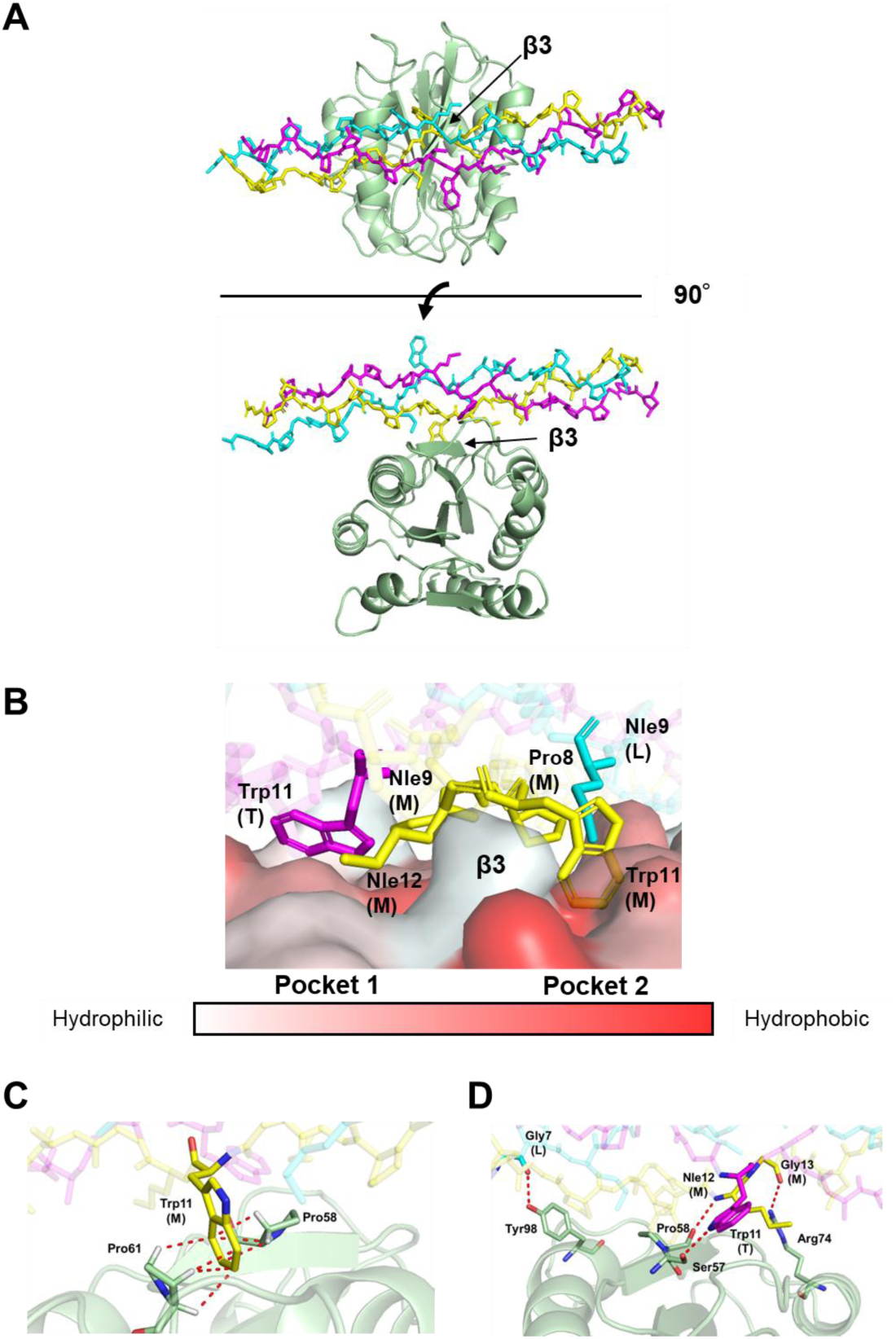
Complex of collagen VII A2 domain with Short-XGWXGA. *A*, Crystal structure of the collagen VII A2–Short-XGWXGA complex. Collagen VII A2 is shown as a cartoon model, and Short-XGWXGA is shown as a stick model. The leading (L), middle (M), and trailing (T) chains of Short-XGWXGA are colored cyan, yellow, and magenta, respectively. The β3 strand of the A2 is labeled. *B*, Two hydrophobic pockets of collagen VII A2 in the complex shown in panel A. Surface hydrophobicity is colored according to the normalized consensus hydrophobicity scale (39). *C*, CH–π interactions (dashed lines) observed in the complex shown in panel A. The hydrogen atoms involved are displayed as sticks in white. *D*, Hydrogen bonds (dashed lines) at the binding interface in the complex shown in panel A.

These structural findings define the molecular basis for A2 recognition of the Met-Gly-Φ motif. Although the Met-Gly-Trp triplet is absent in the sequences of human collagen, a database search for sequences containing the Met-Gly-Φ residue motif identified several candidates including a region within collagen III corresponding to Pep 4, human α1(III) 578–586 (Table 2). Pep 4 was shown to interact with the A2 domain, albeit with substantially lower affinity than Pep 3 (Figure 2A). To examine A2 recognition in a more native collagen context, we performed molecular dynamics (MD) simulations using a triple-helical peptide incorporating a longer collagen III–derived guest sequence, Ac-Gly-Pro-Arg-Gly-Gln-Hyp-Gly-Val-Met-Gly-Phe-Hyp-Gly-Pro-Lys-Gly-Asn-Hyp-Gly-Pro-Hyp-NH₂ (Pep Col3), which encompasses the guest sequence of Pep 4 and corresponds to the human α1(III) 572–586. We constructed a structural model of Pep Col3 in complex with A2 domain, based on the crystal structure of the A2–Short-XGWXGA complex. Energy minimization of the model revealed no steric clashes. The two Phe residues from the middle and trailing chains were well accommodated within hydrophobic pockets 1 and 2, respectively. As shown by the minimum distance (dmin) plot, a 100-ns MD simulation of the complex revealed stable peptide binding overall (Figure 4A). However, the Phe residue from the trailing chain gradually dissociated from Pocket 1, as indicated by an increase in the center-of-mass distance (dcom) at approximately 21.4 ns, whereas the Phe residue from the middle chain remained stably engaged with hydrophobic residues in Pocket 2 throughout the simulation (Figures 4B and 4D, Supplemental Movie 2). By contrast, in an MD simulation of the A2–ShortPep 3 complex, where the guest sequence XGWXGAOGP in Short-XGWXGA was replaced with MGWMGAKGR of Pep 3, Trp residues from the middle and trailing chains remained stably bound within their respective hydrophobic pockets for the entire 100-ns simulation (Figure 4A, 4B and 4C, Supplemental Movie 1). Notably, the Trp residue in Pocket 1 formed a hydrogen bond *via* the Nε1 atom of its indole ring with Ser57 of the A2 domain throughout the simulation, likely contributing to the higher binding affinity compared to Phe, which lacks a side-chain hydrogen bond donor (Figure 4C). Furthermore, the simulation suggested an additional contribution of the electrostatic interaction between Lys15 of the middle chain and Asp77 of the A2 domain (Figure 4E), consistent with the result shown in Figure 2B.

**Figure 4.**
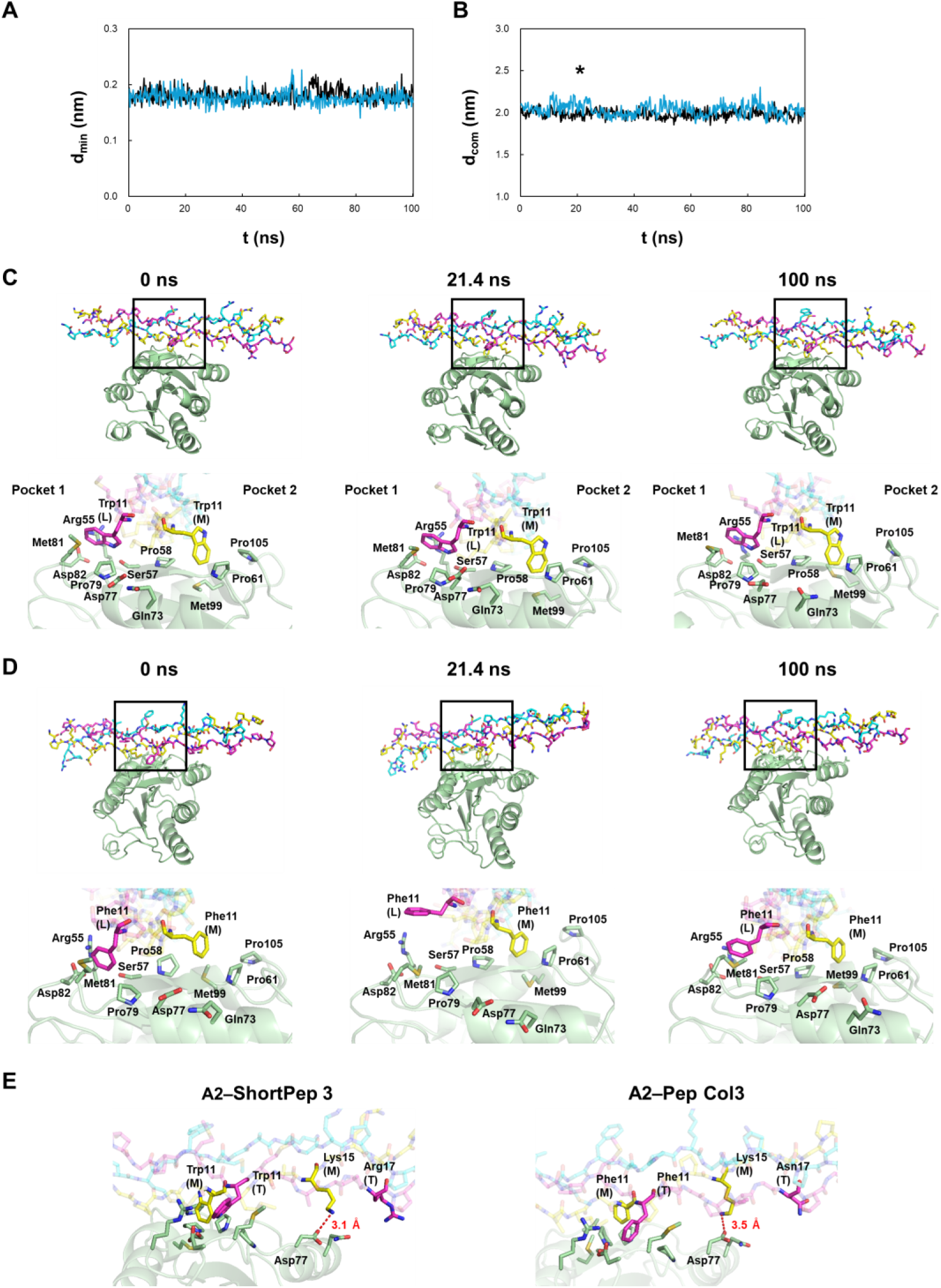
MD simulations of A2 domain–triple-helical peptide interactions. *A*, Time evolution of the minimum distance (dₘᵢₙ) between the A2 domain and the triple-helical peptide. Black line: A2 domain–ShortPep 3 complex; blue line: A2 domain–Pep Col3 complex. *B*, Time evolution of the center-of-mass distance (dcom) between the A2 domain and the triple-helical peptide. The asterisk indicates the time point at which Phe11 of Pep Col3 markedly deviated from Pocket 1. Black line: A2 domain–ShortPep 3 complex; blue line: A2 domain–Pep Col3 complex. *C* and *D*, Snapshots of A2 domain–peptide complexes (ShortPep 3 in C; Pep Col3 in D) at 0, 21.4, and 100 ns. The color scheme for each chain is the same as in Figure 3. Top panels show overall views, with black boxes indicating a binding interface. Bottom panels present close-up views of the boxed regions, showing Trp11 (C) or Phe11 (D) from the middle and trailing chains surrounded by residues of the A2 domain. Pocket 1 and Pocket 2 are labeled. *E*, Snapshots of A2 domain–peptide complexes at 0 ns. Dashed lines indicate electrostatic interactions.

### Binding specificity of the triple-helical peptide for collagen VII A2 domain

The collagen VII A2 domain shares homology with the original VWF A3 domain, which exhibits collagen-binding activity. Similarly, the integrin α2I domain is also homologous to the VWF A3 domain and mediates collagen binding (40). Triple-helical Pep 4 was recognized by collagen VII A2 domain in a metal ion independent manner (Figure S10). To assess the specific recognition of Pep 4, corresponding to human α1(III) 578–586, by the A2 domain, we performed binding assays using peptide-conjugated beads and lysates of *Escherichia coli* (*E. coli*) expressing GST fusion proteins of the A2, VWF A3 domain, and integrin α2I. Pep 5 contains the Arg-Gly-Gln-Hyp-Gly-Val-Met-Gly-Phe, corresponding to α1(III) 572–580, and was used as a VWF A3-binding sequence (41). Pep 6, a peptide containing the Gly-Phe-Hyp-Gly-Met-Arg motif, was used as an integrin α2I-binding sequence (Figure 5A, Table S1) (32). As shown in Figure 5B, Pep 4-immobilized beads specifically pulled down the collagen VII A2 domain from the corresponding *E. coli* lysate, whereas neither VWF A3 nor integrin α2I was retained on the beads. All domains bound to the glutathione-immobilized beads (GST-accept), whereas no binding was observed with mock-coupled beads. As expected, VWF A3 bound to Pep 5, and integrin α2I bound to Pep 6. These results demonstrate that the collagen III-derived Pep 4 interacts specifically with the collagen VII A2 domain. However, binding between the collagen III–derived Pep 5, which shares the overlapping α1(III) 578–580 sequence with Pep 4 (Figure 5A), and the collagen VII A2 domain was not detectable in the pull-down assay.

**Figure 5.**
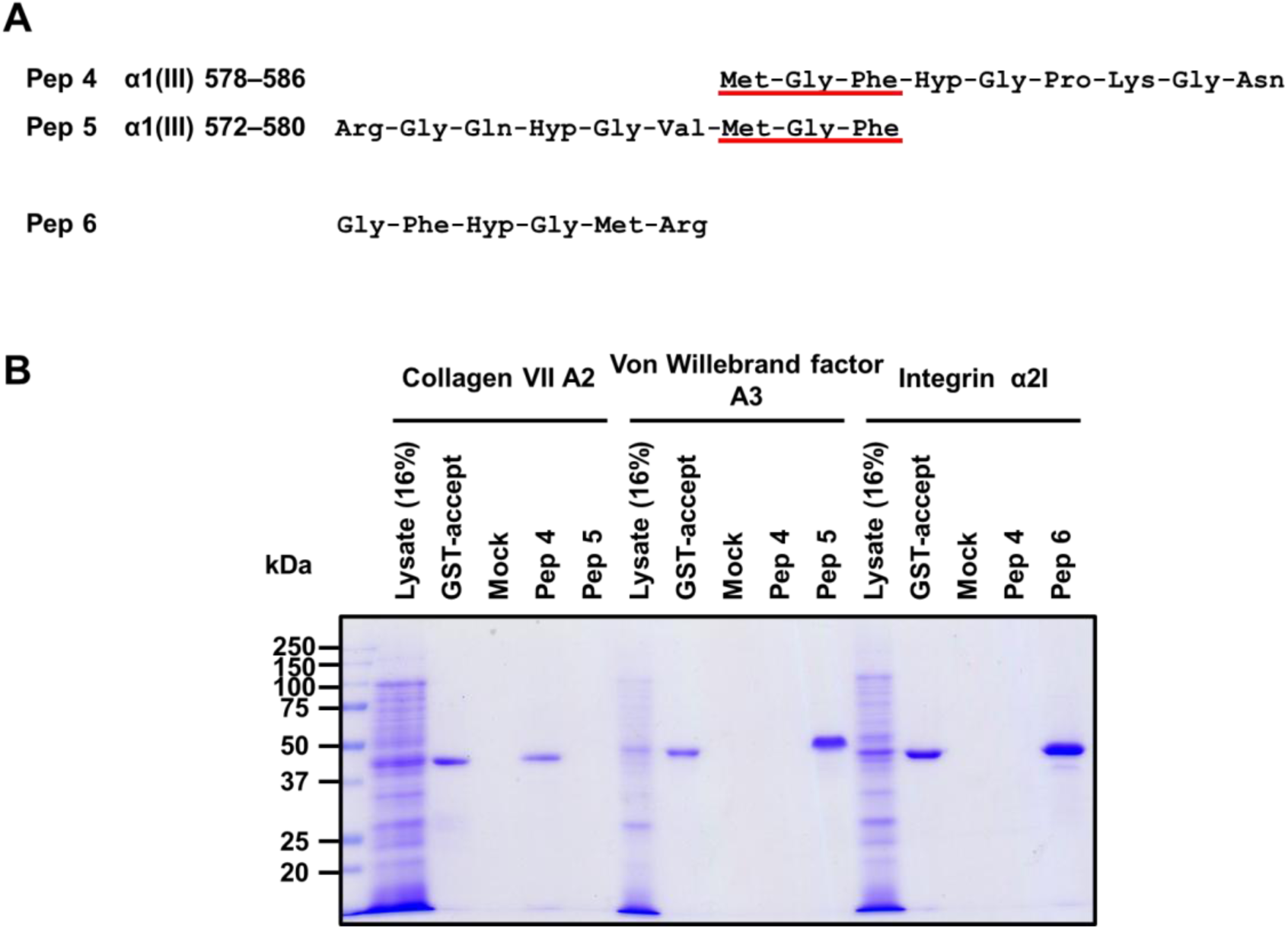
Binding of von Willebrand factor A3-like domains to bead-conjugated peptides. *A*, Guest sequences of the peptides used for peptide-conjugated beads. Overlapping α1(III) 578–580 sequences of Pep 4 and Pep 5 are underlined. *B*, Peptide-conjugated beads were incubated with *E. coli* lysates expressing GST fusion proteins of collagen VII A2, VWF A3, or integrin α2I domains. Beads-bound proteins were separated by 12% SDS-PAGE under a reducing condition and visualized by Coomassie Brilliant Blue (CBB) staining. The molecular sizes of collagen VII A2, VWF A3, and integrin α2I are approximately 48 kDa. GST-accept resin and mock-coupled beads served as positive and negative controls, respectively. Pep 5 and Pep 6 were included as reference peptides for VWF A3 and integrin α2I binding.

To further examine whether the collagen VII A2 domain recognizes sequences overlapping with the VWF A3-binding region, we performed an ELISA (Figure 6). Wells were coated with a C3-Pep 5 polymer, in which the Pep 5 guest sequence, corresponding to α1(III) 572–580, was presented. As a control, a C3 polymer containing Pro-Gly-Pro-Pro-Gly-Pro-Arg-Gly-Pro-Pro as a guest sequence was used. In this assay, the collagen VII A2 domain showed appreciable binding to the C3-Pep 5-coated surface compared with the control C3 polymer. Notably, C3-Pep 5 exhibited an approximately tenfold lower apparent binding affinity for the collagen VII A2 domain compared with C3-Pep 4. This result is consistent with the MD simulation results, which suggested a contributory interaction involving Lys residue in the Pep 4 (Figure 4E). Together, these results show that the collagen VII A2 domain binds to both Pep 4 and Pep 5, which contain the Met-Gly-Phe motif corresponding to α1(III) 578–580, whereas VWF A3 binds to Pep 5 but not to Pep 4. Thus, both the collagen VII A2 domain and VWF A3 recognize an overlapping region within collagen III.

**Figure 6.**
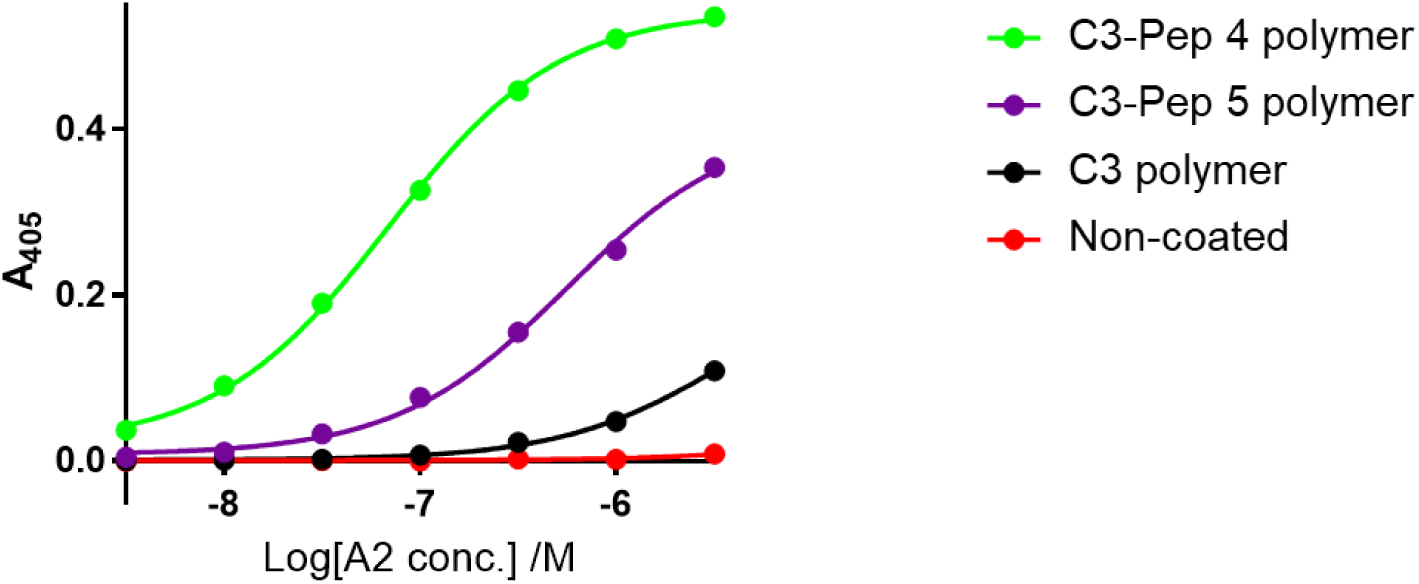
Collagen VII A2 binding to C3-Pep 5 polymer. GST-collagen VII A2 was incubated with ELISA wells coated with peptide polymers (0.5 μg/well) at 25°C. Bound protein was detected using HRP-conjugated anti-GST antibody and ABTS substrate, and absorbance was measured at 405 nm (n = 3, mean ± S.D.).

## Discussion

In this study, we identified specific collagen-like triple-helical sequences recognized by the collagen VII A2 domain and elucidated the molecular basis of this interaction through an X-ray crystallographic analysis. These findings provide new molecular insights into how collagen VII forms arc-shaped anchoring fibrils that integrate into the basement membrane by interacting with mesenchymal collagen triple helices, which traverse through anchoring fibrils.

Previous studies have reported that collagen VII A2 domain binds to collagen I, III, and IV, but the molecular mechanism by which the domain recognizes triple helices has remained unclear (24,26). These interactions with native collagen triple helices have been reported to be relatively weak, which has hindered detailed structural analysis of the A2 domain–triple helix interface (23). To address this, we first performed a yeast two-hybrid screening of triple-helical random peptide library. Consistent with previous reports that collagen VII interacts with collagen *via* the A2 domain, positive colonies were obtained for the A2 domain but not for the A1 domain. The Met-Gly-Φ motif was identified as an A2 domain-binding motif. Particularly, the acquisition of a non-native high-affinity peptide, containing a Met-Gly-Trp motif instead of the native Met-Gly-Phe motif, enabled us to elucidate the molecular basis of the interaction through biochemical analyses and X-ray crystallography. Collectively, the use of a higher-affinity surrogate ligand allowed us to define key features of A2 domain–triple helix recognition that would have been difficult to resolve using native collagen sequences alone.

The triple-helical peptide containing a Met-Gly-Trp motif formed a stable complex with the A2 domain. Notably, the A2 interaction involves all three peptide chains, necessitating assembly of the triple helix. The collagen VII A2 domain and the VWF A3 domain belong to the same Rossmann fold-based superfamily (14). Both domains engage all three chains of the collagen triple helix. Consistent with these shared features, both domains bind sequences containing a Met-Gly-Φ signature, such as Arg-Gly-Gln-Hyp-Gly-Val-Met-Gly-Phe at human α1(III) 572–580, which corresponds to the guest sequence of Pep 5 (41). Thus, collagen VII A2 and VWF A3 share overlapping triple-helical binding motifs confirmed in Figure 6. Nevertheless, collagen VII A2 differs in its recognition of aromatic residues, demonstrating a distinct collagen-binding mode (Figures 3 and 4). In particular, the triple helix is bound in the opposite orientation relative to the VWF A3 complex (Figure S11) (42). These differences highlight that, despite the similarities, collagen VII A2 employs a unique collagen-binding mechanism distinct from that of VWF A3.

To further explore the physiological relevance of this interaction, we combined X-ray crystallographic and MD data. Our MD simulations demonstrated that Trp residues in the high affinity unnatural peptide (Short-MGWMGA) remained stably accommodated within the two hydrophobic pockets of the A2 domain throughout the simulation, forming persistent CH–π and hydrogen-bond interactions. In contrast, when the corresponding Trp residues were replaced by Phe, as found in the native Met-Gly-Phe motif, the side chain of the Phe from the trailing chain gradually dissociated from the Pocket 1 due to the absence of an indole nitrogen capable of hydrogen bonding. These simulations suggest that native collagen sequences bearing the Met-Gly-Phe motif are recognized by A2 more weakly than unnatural Met-Gly-Trp-containing peptides. This simulation was consistent with the result of the *in vitro* binding assay shown in Figure 2A.

In the crystal structure and MD simulation, the Met residues from the leading and middle chains and the aromatic residues from the middle and trailing chains of the Met-Gly-Φ motif interacted with the A2 domain. These residues occupied two distinct hydrophobic pockets that define the binding interface. Consistent with the yeast two-hybrid screening results, Met and aromatic residues at these positions were highly conserved among the identified binding sequences, highlighting their importance for A2 recognition. Collagens I and III, which are components of mesenchymal collagen fibrils, contain the Met-Gly-Phe motif as the Met-Gly-Φ motif. Collagen I forms a hetero-triple helix composed of two α1(I) and one α2(I) chains. In this composition, the α1(I) 580–582 segment (Met-Gly-Phe) corresponds to α2(I) 492–494 (Ile-Gly-Phe). Three possible stagger configurations can occur, α1α1α2, α1α2α1 and α2α1α1. Recent studies suggested that α1α1α2 and α2α1α1 are plausible (43,44). As shown in Figure 7, the α1α1α2 hetero-triple helix configuration has Met in the leading and middle chains and Phe in the middle and trailing chains, thereby fulfilling the binding surface geometry identified by the crystallographic analysis. Even in the other two possible staggering isomers of collagen I, the A2 domain-binding surface seems to be preserved, although each Met residue is replaced by Ile. Similarly, collagen III, which forms a homotrimer, possesses a Met-Gly-Phe sequence at α1(III) 578–580 that satisfies the same geometric surface structure for A2 recognition, and the interactions with the sequences were actually confirmed in this study (Figures 2, 5 and 6). In addition to the hydrophobic interactions mediated by the Met-Gly-Φ motif, the MD simulation of Pep Col3 suggested an additional electrostatic interaction between Lys15 of the middle chain and Asp77 of the A2 domain (Figure 4E). As collagen I and III sequences also contain Lys residues at the corresponding position, this auxiliary interaction is preserved (Figure 7). Collagen IV, which exists at the DEJ as α1α1α2 and α5α5α6 heterotrimers (45), also contains Met-Gly-Phe motifs, raising the possibility that the collagen VII A2 domain may also recognize these sequences. However, in the mature DEJ, collagen VII is thought to interact mainly with the globular NC1 domains of collagen IV rather than its triple-helical region (20). Although collagen IV carries a Met-Gly-Phe motif, the stable interaction appears to be mediated by its globular NC1 domains.

**Figure 7.**
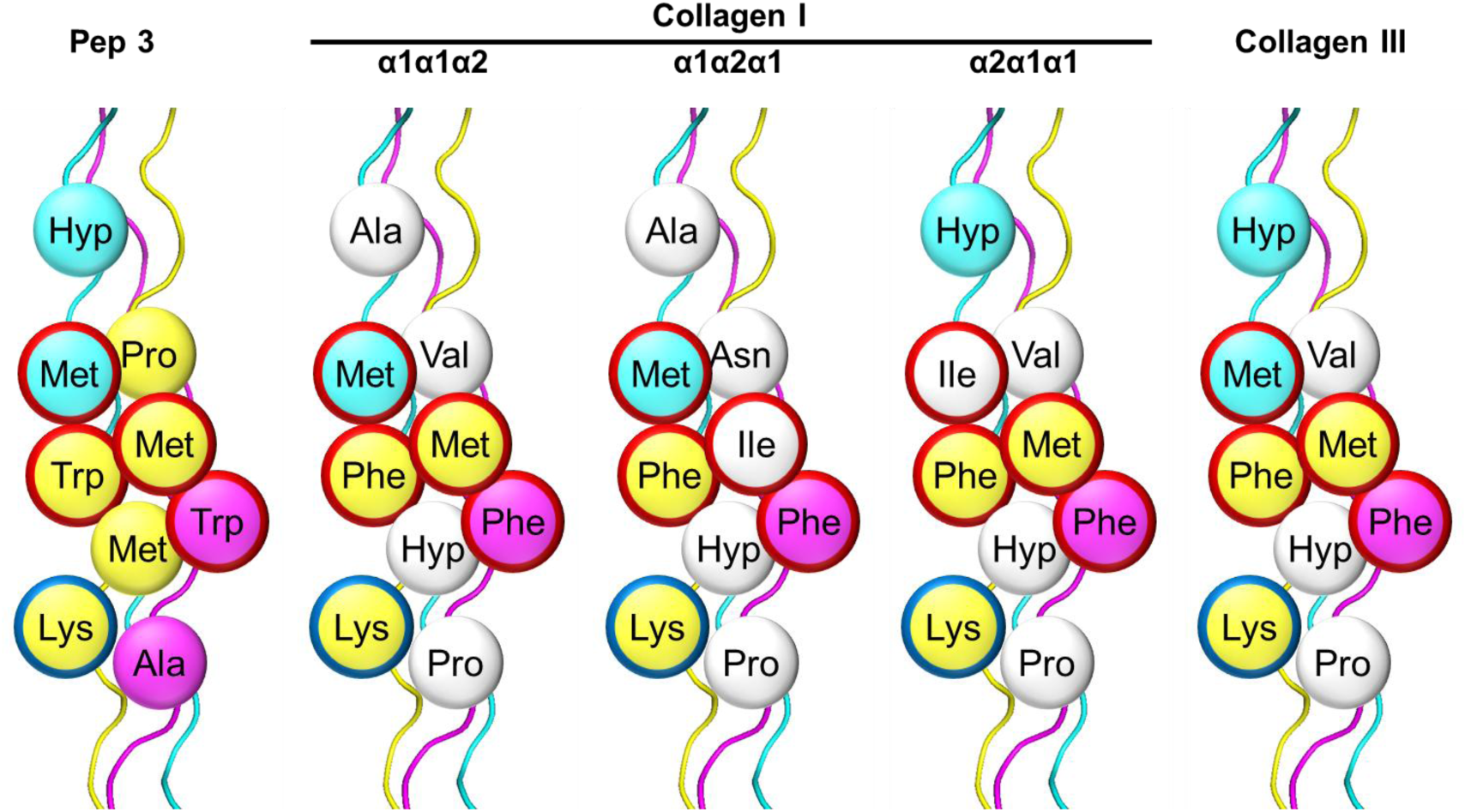
Modeling of amino acid arrangement of the collagen triple helix. Models were constructed based on the triple-helical peptide corresponding to Short-XGWXGA peptide extracted from the co-crystal structure determined in this study. The leading, middle, and trailing chains are colored cyan, yellow, and magenta, respectively. The modeled structures include Pep 3, three heterotrimeric configurations of collagen I, and a homotrimeric configuration of collagen III. In the comparison between Pep 3 and collagen sequences, Trp residues in the peptide were treated as representative residues for aromatic residues. Residues that differ from those in Pep 3 are shown in white. Amino acid residues at the positions of the Met-Gly-Φ motif whose side-chains are accommodated into the two pockets of the A2 domain are outlined in red. Lys residues for electrostatic interactions are outlined in blue.

Based on these observations, we propose two models for anchoring fibril integration into the DEJ. In one model (Figure 8A), anchoring fibrils first interact with nearby collagen fibrils through transient A2-mediated recognition of the Met-Gly-Phe motif, thereby promoting the positioning of collagen fibrils inside the arc-shaped structure of anchoring fibrils, prior to stable attachment to the basement membrane. In the alternative model (Figure 8B), one end of the anchoring fibril initially binds to the components of the basement membrane, such as collagen IV and laminin 332, while the opposite end contacts collagen fibrils *via* the A2 domain. Anchoring fibrils recruit collagen fibrils and enclose them within their arcs, with both ends of the anchoring fibrils ultimately secured to the basement membrane. Notably, the Met-Gly-Phe motifs on collagen I and III, which are recognized by the collagen VII A2 domain, are exposed to the surface of the fibrils and remain accessible (46,47). The Met-Gly-Phe motif is also recognized by other collagen-binding proteins, namely VWF A3, discoidin domain receptors (DDRs), and secreted protein acidic and rich in cysteine (SPARC), highlighting its functional importance in mediating interactions (47). Previous biochemical studies have shown that the interaction of the collagen VII NC1 domain with the collagen IV NC1 domain or with laminin 332 is approximately 200-fold stronger than its interaction with collagen I, based on differences in dissociation constants (Kd) (23). Although the affinity of the A2 domain for mesenchymal collagen triple helices is comparatively weak, clustering of A2 domains at the anchoring fibril termini is expected to enhance overall interaction through avidity effects, while the flexible hinge region of collagen VII permits the arched conformation (15). In a physiological context, where multiple collagen VII molecules and collagen triple helices are spatially organized within anchoring fibrils, such multivalency is likely to stabilize these otherwise transient interactions and render them sufficient to support fibril assembly and function. Anchoring fibrils are ultimately secured to the basement membrane probably through the firm binding of collagen VII NC1 domain, which contains the A2 domain, to collagen IV globular domains and laminin 332 (20–23,27), ensuring a robust connection. Since A2-mediated interactions with mesenchymal collagen triple helices are not observed in mature anchoring fibrils, the transient interactions between the A2 domain and collagen I/III likely function at earlier assembly stages, facilitating the initial capture and the recruitment of fibrillar collagens into the arc-shaped anchoring fibrils. These models are consistent with the biology of collagen VII, which is secreted by keratinocytes and fibroblasts, and is assembled after the basement membrane and collagen fibrils have been established during skin development (48–52). In this configuration, the arched anchoring fibrils enclose collagen fibrils in a manner reminiscent of polyrotaxanes, where a ring encircles an axle (53,54). This architecture combines flexibility with mechanical strength, suggesting that the arched fibril structure stabilizes the DEJ similarly to how polyrotaxane structures distribute force and resist deformation (55).

**Figure 8.**
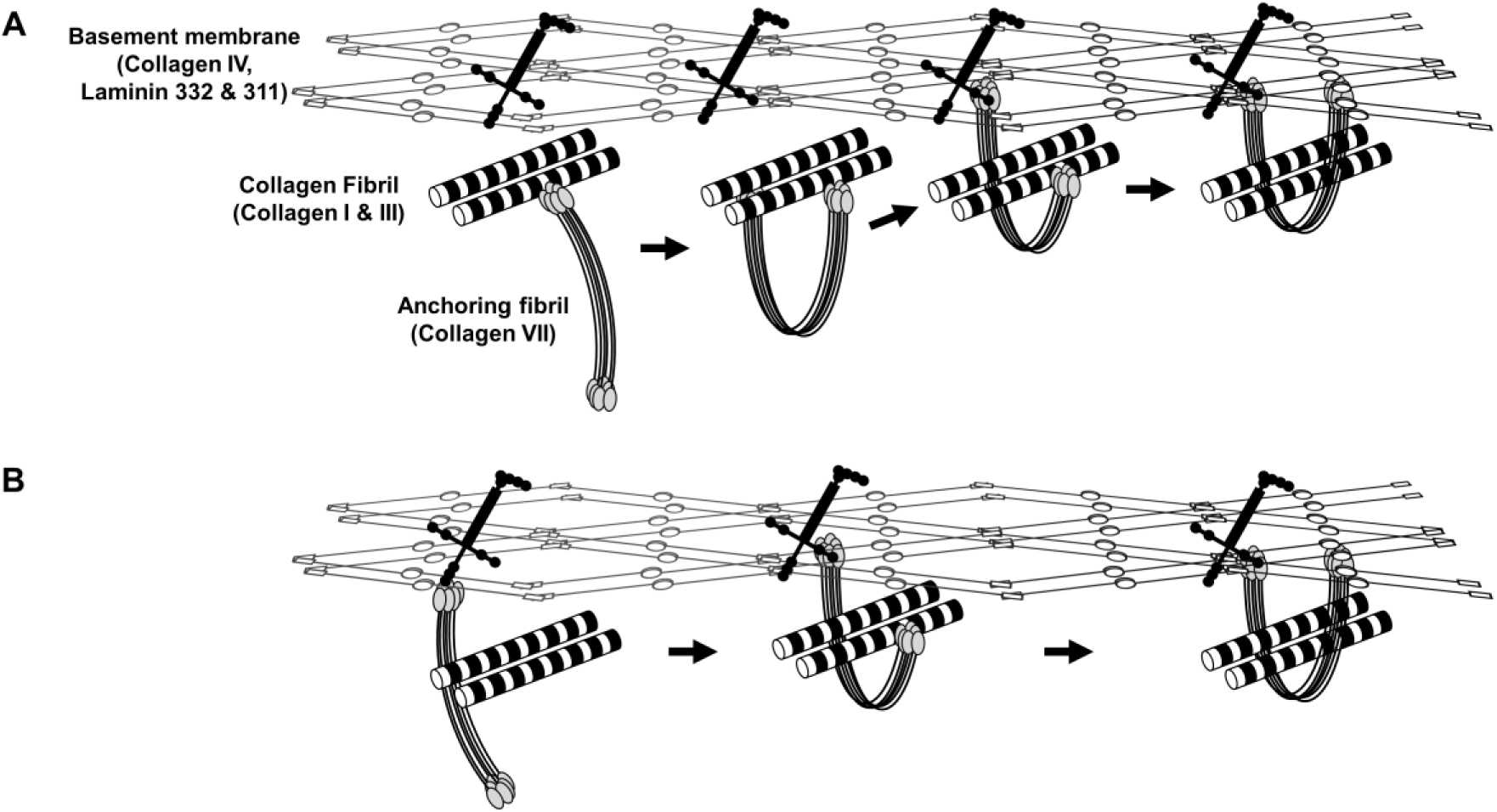
Proposed assembly models of anchoring fibrils integrating the basement membrane. *A*, Anchoring fibrils interact with collagen fibrils through transient A2-mediated recognition before attaching to the basement membrane. *B*, One end of the fibril initially binds the basement membrane while the other engages collagen fibrils.

These molecular insights also provide a basis for connecting structural understanding to disease. In EBA, the A2 domain serves as the major autoantigen, underlining the importance of clarifying its role in maintaining skin stability. By elucidating the molecular basis of the A2 domain interactions with the collagen triple helix, our study has advanced this understanding. Further studies leveraging these structural insights may guide the rational design of biomaterials or therapeutic strategies to restore or enhance anchoring fibril function in EBA.

## Experimental procedures

### Plasmid construction

Plasmids encoding the human collagen VII A1 domain (Val29–Pro227; GenBank accession number AAA58965), the human VWF A3 domain (Ala1683–Gly1874; GenBank accession number AAB59458), and the human collagen VII A2 domain (Val1047–Ser1237; GenBank accession number AAA58965) were synthesized by FASMAC (Kanagawa, Japan). The plasmid encoding the human integrin α2I domain was obtained as previously described (32). Each insert was cloned into the *EcoR*I and *Sal*I sites of the pUCFa vector. Subsequently, the inserts were subcloned into the pGEX-5X-1 vector using the same restriction enzymes, and the resulting construct was transformed into *E. coli* strain HST08.

To generate a GAL4-BD-fusion construct, human collagen VII A1 domain and A2 domain sequences were ligated into the pGBKT7 vector at the *EcoR*I and *Sal*I sites. The resulting plasmids were propagated in *E. coli* HST08 cells.

To construct the A2 domain-expression plasmid for crystallization, the A2 domain coding region was amplified by PCR from the A2-pGEX-5X-1 template (primers: 5’-GTTTGTCCAAGAGGATTAGCTGATG-3’ and 5’-TTATGAAGCTTGACATAAAGCAGTGG-3’). The amplified fragment was assembled into a pET32b backbone prepared by inverse PCR (primers: 5’-TTTATGTCAAGCTTCATAAATGAAAGAAACCGCTGCTGCTAAATTCG-3’ and 5’-GCTAATCCTCTTGGACAAACACCAGAACCGCGTGGCAC-3’) using the NEBuilder® HiFi DNA Assembly Master Mix (New England Biolabs, Ipswich, MA, USA). The resulting plasmid was verified by sequencing and transformed into *E. coli* strain C3029 for protein expression.

### Y2H screening

Yeast strains Y187 and Y2HGold (TaKaRa, Shiga, Japan) were utilized as host cells expressing GAL4-AD-fused triple-helical peptides and GAL4-BD-fused human collagen VII A1 domain and A2 domain, respectively. Transformation of yeast with plasmid DNA was conducted following the manufacturer’s instructions. Screening for triple-helical peptides that interact with the A1 domain and A2 domain were carried out as previously described (31). Briefly, the Y2HGold strain harboring the A1 and A2 constructs were cultured in SD/-Trp broth at 30°C until reaching an optical density at 600 nm (OD600) of 0.5–0.6. The cultures were then mixed with Y187 expressing AD-fused triple-helical random peptides. The mixtures were filtered through a 0.45 µm nylon membrane (GE Healthcare Life Sciences, Pittsburgh, PA, USA), which were subsequently incubated on yeast peptone dextrose adenine (YPDA) plate overnight at 30°C. Following incubation, yeast cells were recovered in saline and plated onto SD/-Leu/-Trp/-His (triple dropout; TDO) selection plates supplemented with 200 ng/mL aureobasidin. Interaction assays were performed by ligating ssDNA sequences into the pGADT7-AD vector. Seq *a* was used as previously reported (31). For Clones 1, 4, and 7, plasmids were extracted from diploid yeast hits using the Zymoprep^TM^ Yeast Plasmid Miniprep I kit (Zymo Research, Irvine, CA, USA), transformed into HST08, and cultured in LB with ampicillin. The plasmids were then amplified, extracted, and transformed into Y187, which was cultured on SD/-Leu plates. The experimental procedures followed the previously reported method. TDO plates were used instead of quadruple dropout (SD/-Leu/-Trp/-His/-Ade) plates (31).

### DNA Sequencing

To identify the peptide sequences of selected clones, individual colonies grown on TDO plates were picked and used as templates for PCR amplification with primers (32). PCR products were purified using the MagExtractor^TM^ PCR & Gel Clean up Kit (TOYOBO, Tokyo, Japan).

Purified amplicons were subjected to Sanger sequencing, which was performed by FASMAC.

### Database search

BLAST searches against the UniProt database (https://www.uniprot.org/blast) were performed using Met-Gly-Trp, Met-Gly-Phe and Met-Gly-Tyr as query sequences to identify corresponding regions in human collagen proteins.

### Peptide synthesis and characterization

To ensure that the peptides adopt a triple helix at 25°C, the thermal stability of candidate sequences was predicted using SCEPTTr (33). C3 constructs were prepared according to previously reported methods (37). Peptides were synthesized manually by the standard 9-fluorenylmethoxycarbonyl (Fmoc)-based solid-phase peptide synthesis strategy. C3-Pep 4 and C3 were constructed on 2-chlorotrityl chloride resin (Peptide Institute, Osaka, Japan), whereas other peptides were synthesized on Rink amide resin (Novabiochem, San Diego, CA, USA). Coupling reactions were carried out at room temperature for 1.5 hours using 3–5 equivalents of Fmoc-protected amino acids activated with 1-hydroxybenzotriazole (HOBt) and *N,N*′-diisopropylcarbodiimide (DIC) in *N,N*-dimethylformamide (DMF). Fmoc group removal was accomplished using 20% (v/v) piperidine in DMF for 20 minutes. Cleavage from the resin and side-chain deprotection were performed by treating the peptide-resin with trifluoroacetic acid (TFA)/H₂O/*m*-cresol/thioanisole/3,6-dioxa-1,8-octanedithiol (80/5/5/5/5, v/v) for 3 hours at room temperature.

Crude peptides were purified by reversed-phase high-performance liquid chromatography (RP-HPLC) using a Cosmosil 5C18 AR-II column (20 mm I.D. × 250 mm; Nacalai Tesque, Kyoto, Japan). The purified products were characterized by electrospray ionization time-of-flight (ESI-TOF) mass spectrometry (Compact; Bruker, or TripleTOF® 4600; AB Sciex). Observed molecular weights matched the theoretical values for all peptides.

Concentrations of soluble peptides were determined by ultraviolet absorbance at 280 nm, based on the molar extinction coefficients of Tyr and Trp residues (56). C3-Pep 4, C3-Pep 5, and C3 were annealed in 0.05% TFA/H₂O and polymerized as described previously (37). To form disulfide cross-links, dimethyl sulfoxide (DMSO) was added to a final concentration of 10% (v/v), and the peptide solution was incubated at 4°C for one week.

### CD spectrometry of the synthetic peptide

CD measurements were performed using a J-820 spectropolarimeter (JASCO, Tokyo, Japan) equipped with a Peltier temperature controller and a 0.5-mm path-length quartz cuvette. The system was connected to a data processing unit for signal averaging. Peptides dissolved in distilled water were heated at 95°C for 5 minutes, cooled at room temperature for 10 minutes, and stored at 4°C overnight to allow triple-helix formation. CD spectra were recorded from 200 to 260 nm in continuous scanning mode, and results were expressed as mean residue ellipticity ([θ]mrw). The triple-helix melting curve was obtained by monitoring the change in [θ]225 as the temperature increased from 4 to 85°C at a rate of 18°C per hour.

### Expression and purification of GST-collagen VII A2 domain

An *E. coli* BL21(DE3) strain harboring a plasmid encoding the GST-tagged human collagen VII A2 domain was cultured in lysogeny broth (LB) supplemented with 100 μg/mL ampicillin at 37°C with shaking. When the OD600 reached between 0.7 and 1.2, protein expression was induced by the addition of 0.1 mM isopropyl-*β*-D-1-thiogalactopyranoside (IPTG), followed by incubation at 37°C for 4 hours. Cells were harvested by centrifugation at 1,660 × *g*, for 10 minutes at 4°C, washed once with phosphate-buffered saline (PBS), and centrifuged again under the same conditions. The resulting pellet was resuspended with 1 mg/mL lysozyme in lysis buffer containing 50 mM Tris-HCl (pH 8.0), 150 mM NaCl, 5 mM EDTA, and 30% (v/v) glycerol. Protease inhibitor cocktail and 0.2% NP-40 Alternative (EMD Millipore, Burlington, MA, USA) were added, and the suspension was rotated for 15 minutes at 4°C. Genomic DNA was fragmented by sonication, and cellular debris was removed by centrifugation. The clarified lysate was incubated with GST-accept resin (Nacalai Tesque) for 1 hour at 4°C. After binding, the resin was sequentially washed with PBS, 0.4 M NaCl in PBS, and PBS. The bound protein was eluted using buffer containing 50 mM Tris-HCl (pH 8.0) and 10 mM reduced glutathione. The eluate was dialyzed against PBS, flash-frozen in liquid nitrogen, and stored at -24°C until use. Protein concentration was determined by the Bradford assay using bovine serum albumin (BSA) as a standard. The purity of the GST-collagen VII A2 domain was confirmed by SDS-PAGE on a 12% polyacrylamide gel (Figure S12).

### ELISA

Flat-bottom 96-well plates (161093, Nunc, Kamstrup, Denmark) were coated with 50 μL of 10 μg/mL peptide polymers at room temperature for 1 hour. After coating, the solution was removed, and wells were blocked with 100 μL of 2 mg/mL skim milk in ELISA buffer (0.05% Tween 20 in PBS) at room temperature for 1 hour. The wells were washed with ELISA buffer three times, and 50 μL of PBS containing 0.1 mg/mL skim milk and the GST-collagen VII A2 domain was added to each well. For the ELISA shown in Figure 6, the GST-collagen VII A2 domain was incubated in PBS containing 0.05% Tween 20, 0.1 mg/mL skim milk, and 0.4 M NaCl. After incubation at 24°C for 90 minutes, wells were washed with ELISA buffer five times. Horseradish peroxidase (HRP)-conjugated anti-GST antibody (GE Healthcare Life Sciences) diluted 1:3000 in ELISA buffer was added and incubated at 4°C for 45 minutes. After washing the wells three times, 50 μL of 0.5 mg/mL 2,2′-azinobis(3-ethylbenzthiazoline-6-sulfonic acid) diammonium salt (Fujifilm Wako Pure Chemical Industries, Osaka, Japan) in citrate-phosphate buffer containing 0.1 M citrate, 0.2 M Na2HPO4 (pH 5.0) was added to each well. Plates were incubated at 37°C for 30 minutes and absorbance at 405 nm was measured.

For the competitive assay, the GST-collagen VII A2 domain was preincubated at a constant concentration (5 nM or 10 nM) with various concentrations of soluble peptides.

For the metal ion-dependency assay, the GST-collagen VII A2 domain was diluted in either 2 mM MgCl₂ or 5 mM EDTA.

### Crystallization

Cells were cultured in LB medium containing ampicillin at 37°C until the OD660 reached 0.6, and protein expression was induced with 0.2 mM IPTG at 20°C overnight. Cells were harvested by centrifugation and resuspended in lysis buffer [20 mM Tris-HCl (pH 8.0), 1 M NaCl]. The suspension was treated with lysozyme for 15 min, followed by Triton X-100 for 15 min. Cells were disrupted by sonication, and insoluble material was removed by centrifugation at 24,000 × *g* for 1 h at 4°C. The supernatant was loaded onto a Ni Sepharose FF column (Cytiva, Marlborough, MA, USA) equilibrated with buffer A [20 mM Tris-HCl (pH 8.0), 1 M NaCl]. The column was washed with buffer A and buffer B [20 mM Tris-HCl (pH 8.0), 150 mM NaCl, 20 mM imidazole], and bound proteins were eluted with buffer C [20 mM Tris-HCl (pH 8.0), 150 mM NaCl, 500 mM imidazole]. The eluate was dialyzed against buffer D [20 mM Tris-HCl (pH 8.0), 150 mM NaCl] in the presence of thrombin to remove the His tag. The digestion mixture was re-applied to the Ni Sepharose FF column, and the flow-through containing tag-free A2 domain was collected. Final purification was performed by size-exclusion chromatography on a Superdex 75 16/600 column (Cytiva) equilibrated with buffer D.

Purified A2 domain was concentrated to 5 mg/mL (243 μM) and mixed with Short-XGWXGA peptide (291 μM) at a molar ratio of 1:1.2. Crystallization screening was performed at 4°C by the sitting-drop vapor diffusion method using Crystal Screen I and II (Hampton Research). Plate-like crystals appeared under several conditions. Diffraction-quality crystals were obtained in reservoir solution containing 26% (w/v) PEG 4000, 100 mM sodium acetate, and 100 mM ammonium sulfate at pH 4.8.

### Data collection, structure determination, and refinement

Crystals of the collagen VII A2–Short-XGWXGA complex were picked up from the crystallization drop using a nylon loop, and flash cooled in liquid nitrogen. X-ray diffraction data were collected at beamline BL45XU of the synchrotron radiation facility SPring-8 using an EIGER X 16M detector at 100 K. The diffraction images were processed with XDS (57) and scaled with Aimless in the CCP4 suite (58). Initial phases were obtained by molecular replacement using PHASER, as implemented in the PHENIX package (59). A predicted model of the A2–triple-helical peptide complex, generated by the AlphaFold Server (60), was used as the search model. The peptide sequence used for modeling was Ac-(Gly-Pro-Pro)₂-Gly-Pro-Gly-Trp-Leu-Gly-Ala-Pro-(Gly-Pro-Pro)₂-NH₂, in which norleucine and hydroxyproline residues in the Short-XGWXGA were substituted with leucine and proline, respectively. The molecular replacement solution revealed two complexes per asymmetric unit. Model building, including the placement of water molecules and the substitution of Nle and Hyp residues, was performed using Coot (61). Iterative rounds of refinement with *phenix.refine* and manual adjustments in Coot (61) yielded the final structure at 1.47 Å resolution, with *R*work and *R*free values of 0.176 and 0.217, respectively. Model validation was performed with MolProbity (62). Data collection and refinement statistics are summarized in Table S3.

### Molecular dynamics simulations

All simulations were performed with GROMACS 2025.2 (http://www.gromacs.org/) (63). The initial coordinates were based on the crystal structure determined in this study of the collagen VII A2 domain in complex with a short collagen peptide (A2–Short-XGWXGA). The guest sequence OGPOGPXGWXGAOGP in the peptide was replaced in Coot (61) with either Pep 3 (OGPOGPMGWMGAKGR) or the human α1(III) 572–586 (RGQOGVMGFOGPKGR).

Topologies were generated with AMBER99SB-ILDNP and TIP3P water. Each system was placed in a cubic box with at least 1.2 nm buffer from the box edge, solvated with TIP3P water, and neutralized by replacing solvent molecules with Cl⁻ counter-ions. Energy minimization was performed using the steepest descent algorithm (50,000 steps; convergence threshold emtol = 100 kJ mol⁻¹ nm⁻¹). Equilibration consisted of (i) a 100 ps NVT run with heavy-atom position restraints on the solute, (ii) a 500 ps unrestrained NVT run, and (iii) a 500 ps NPT run. Temperature was maintained at 300 K using the V-rescale thermostat (τ_T = 0.1 ps), and pressure was controlled at 1 bar using the Parrinello–Rahman barostat (τ_P = 2.0 ps; isotropic coupling; compressibility = 4.5×10⁻⁵ bar⁻¹). Long-range electrostatic interactions were calculated with the Particle Mesh Ewald (PME) method, and the Verlet cutoff scheme was applied with real-space cutoffs of 1.0 nm for both Coulomb and Lennard–Jones interactions. All bonds involving hydrogen atoms were constrained with the Linear Constraint Solver (LINCS) algorithm. A 2 fs integration step was used. Periodic boundary conditions were applied in all three dimensions.

Production MD simulations were run for 100 ns for each system, saving coordinates and velocities every 20 ps and energies/logs every 2 ps. To ensure reproducibility and sufficient sampling, three independent simulations were performed for each peptide system, starting from identical coordinates but with different random seeds for initial velocity generation. The corresponding movies can be seen in Supplemental Movies.

### Distance analyses

Distances between the A2 domain and the triple-helical collagen peptide (three chains combined into one group) were computed using GROMACS 2025.2 (http://www.gromacs.org/) (63). The minimum interatomic distance (dmin) was defined as the shortest atom–atom distance between the A2 domain and collagen throughout the trajectories. The center-of-mass distance (dcom) was calculated as the Euclidean distance between the centers of mass of the A2 domain and the combined collagen chains. Predefined index groups were used in all calculations, and time series of dmin and dcom were analyzed to evaluate the stability and binding mode of the complexes.

### Solid-phase binding assay

Peptides were immobilized onto cyanogen bromide (CNBr)-activated sepharose 4B (Cytiva) following the manufacturer’s protocol. The peptide-conjugated beads were refolded at 4°C overnight before the binding assay. *E. coli* lysates expressing the GST-collagen VII A2 domain, the GST-VWF A3 domain, and the GST-integrin α2I domain were prepared as described above, except that the induction was carried out at 25°C instead of 37°C.

For the binding assay, 220 µL of each lysate was mixed with 20-µL bed of the affinity beads. The reaction was carried out at 4°C for 1 hour with gentle rotation. The supernatant was discarded, and the beads were washed three times with PBS. Proteins retained on the beads were eluted by adding 20 μL of 2 × sample buffer containing 50 mM Tris–HCl (pH 6.7), 2% SDS, 10% glycerol, 0.002% bromophenol blue, followed by the addition of 100 mM dithiothreitol (DTT), and heating at 95°C for 5 minutes. Samples were separated by 12% SDS-PAGE and visualized by CBB-staining.

## Data availability

The structure data of human collagen VII A2 domain in complex with homo-trimeric collagen model peptide Short-XGWXGA have been deposited in the Protein Data Bank (https://www.rcsb.org) (PDB ID: 9X02).

## Supporting information

This article contains supporting information (31,32,33,37,38,42,64).

