## Supporting Information for "Molecular basis of collagen triple helix recognition by VWF A-like domain 2 of collagen VII: Implications for interlaced anchoring fibril formation"

**Table of contents**

1. The two-hybrid interaction of the AD-fusion peptides with the triple-helical random peptide library and the BD-fusion collagen VII A2 domain, HSP47, and PEDF. (Figure S1)	S-2
2. Amino acid sequences of synthesized peptides. (Table S1)	S-3–S-4
3. Reverse-phase high-performance liquid chromatography (RP-HPLC) profiles of synthesized peptides. (Figure S2)	S-5–S-6
4. Mass spectrometry (MS) charts of the synthetic peptides. (Figure S3)	S-7–S-15
5. Circular dichroism (CD) profiles of peptides at 4°C (solid line), 25°C (dashed line), and 85°C (dash-dotted line). (Figure S4)	S-16–S-17
6. CD spectra recorded at 225 nm from 4 to 85°C. (Figure S5)	S-18
7. Predicted melting temperatures (*T*_m_) of triple-helical peptides containing Trp at Xaa3. (Table S2)	S-18
8. Collagen VII A2 binding increased with the proportion of the C3-Pep 4 in the peptide polymer. (Figure S6)	S-19
9. Data collection and refinement statistics. (Table S3)	S-20
10. Co-crystal of collagen VII A2 and Short-XGWXGA. (Figure S7)	S-21
11. Collagen VII A2 domain recognizes Nle-containing triple-helical peptide. (Figure S8)	S-22
12. Structural comparison of collagen VII A2 between the apo and complex forms. (Figure S9)	S-23
13. Collagen VII A2 recognizes the collagen *triple-helical* peptides in a metal ion-independent manner. (Figure S10)	S-24
14. Comparison of collagen-binding modes of collagen VII A2 and VWF A3. (Figure S11)	S-25
15. SDS-PAGE analysis of purified GST-collagen VII A2 domain. (Figure S12)	S-26
16. References	S-26–S-27

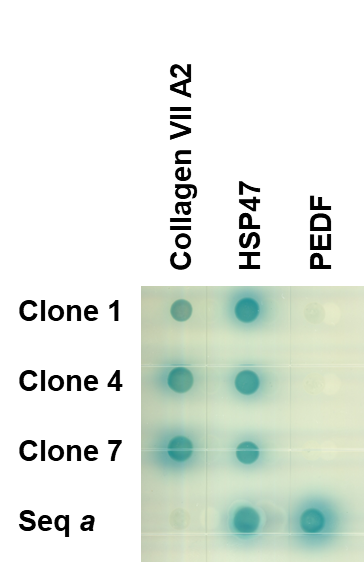

**Figure S1. The two-hybrid interaction of the AD-fusion peptides with the triple-helical random peptide library and the BD-fusion collagen VII A2 domain, HSP47, and PEDF.** The cells were cultured at 25°C for 4 days. The interaction allows yeast cells to survive and turn blue by degrading X-α-Gal. Seq *a* is Lys-Gly-His-Arg-Gly-Phe-Ser-Gly-Leu (1).

**Table S1. Amino acid sequences of synthesized peptides.**

| **Name** | **Sequence** |
| --- | --- |
| Pep 0  Pep 1  Pep 2  Pep 3  Pep 4  Pep 5  Pep 6  C3-Pep 4  C3-Pep 5  C3 | H-(Pro-Hyp-Gly)_6_-Pro-Arg-Gly-(Pro-Hyp-Gly)_5_-Pro-Tyr-NH_2_  H-(Pro-Hyp-Gly)_4_-Pro-Met-Gly-Phe-Gln-Gly-Leu-Arg-Gly-Asp-Hyp-Gly-(Pro-Hyp-Gly)_4_-Pro-Tyr-NH_2_  H-(Pro-Hyp-Gly)_4_-Pro-Met-Gly-Phe-Met-Gly-Ala-Met-Gly-Ala-Hyp-Gly-(Pro-Hyp-Gly)_4_-Pro-Tyr-NH_2_  H-(Pro-Hyp-Gly)_4_-Pro-Met-Gly-Trp-Met-Gly-Ala-Lys-Gly-Arg-Hyp-Gly-(Pro-Hyp-Gly)_4_-Pro-Tyr-NH_2_  H-(Pro-Hyp-Gly)_4_-Pro-Met-Gly-Phe-Hyp-Gly-Pro-Lys-Gly-Asn-Hyp-Gly-(Pro-Hyp-Gly)_4_-Pro-Tyr-NH2  H-(Pro-Hyp-Gly)_5_-Pro-Arg-Gly-Gln-Hyp-Gly-Val-Met-Gly-Phe-Hyp-Gly-(Pro-Hyp-Gly)_5_-Pro-NH_2_  H-(Pro-Hyp-Gly)_5_-Phe-Hyp-Gly-Met-Arg-Gly-(Pro-Hyp-Gly)_5_-Pro-Tyr-NH_2_ (ref. 2)  H-Cys-Cys-Cys-(Pro-Hyp-Gly)_5_-Pro-Met-Gly-Phe-Hyp-Gly-Pro-Lys-Gly-Asn-Pro-Gly-(Pro-Hyp-Gly)_5_-Cys-Cys-Cys-OH  H-Cys-Cys-Cys-(Pro-Hyp-Gly)_5_-Pro-Arg-Gly-Gln-Hyp-Gly-Val-Met-Gly-Phe-Hyp-Gly-(Pro-Hyp-Gly)_5_-Cys-Cys-Cys-OH  H-Cys-Cys-Cys-(Pro-Hyp-Gly)_5_-(Pro-Pro-Gly)_2_-Pro-Arg-Gly-Pro-Pro-Gly-(Pro-Hyp-Gly)_5_-Cys-Cys-Cys-OH (ref. 3) |

(^D^Pro indicates D-Pro.)

Table S1. (*Continued*).

| **Name** | **Sequence** |
| --- | --- |
| ss-Pep 4  Pep 3-X  Pep 3-1  Pep 3-2  Pep 3-3  Pep 3-4  Pep 3-5  Pep 3-6  Pep-XGWXGA  Short-XGWXGA | H-(Pro-Hyp-Gly)_2_-^D^Pro-Hyp-Gly-Pro-Hyp-Gly-Pro-Met-Gly-Phe-Hyp-Gly-Pro-Lys-Gly-Asn-Hyp-Gly-(Pro-Hyp-Gly)_2_-^D^Pro-Hyp-Gly-Pro-Hyp-Gly-Pro-Tyr-NH_2_  H-(Pro-Hyp-Gly)_4_-Pro-Nle-Gly-Trp-Nle-Gly-Ala-Lys-Gly-Arg-Hyp-Gly-(Pro-Hyp-Gly)_4_-Pro-Tyr-NH_2_  H-(Pro-Hyp-Gly)_4_-Pro-Hyp-Gly-Trp-Met-Gly-Ala-Lys-Gly-Arg-Hyp-Gly-(Pro-Hyp-Gly)_4_-Pro-Tyr-NH_2_  H-(Pro-Hyp-Gly)_4_-Pro-Met-Gly-Pro-Met-Gly-Ala-Lys-Gly-Arg-Hyp-Gly-(Pro-Hyp-Gly)_4_-Pro-Tyr-NH_2_  H-(Pro-Hyp-Gly)_4_-Pro-Met-Gly-Trp-Hyp-Gly-Ala-Lys-Gly-Arg-Hyp-Gly-(Pro-Hyp-Gly)_4_-Pro-Tyr-NH_2_  H-(Pro-Hyp-Gly)_4_-Pro-Met-Gly-Trp-Met-Gly-Pro-Lys-Gly-Arg-Hyp-Gly-(Pro-Hyp-Gly)_4_-Pro-Tyr-NH_2_  H-(Pro-Hyp-Gly)_4_-Pro-Met-Gly-Trp-Met-Gly-Ala-Hyp-Gly-Arg-Hyp-Gly-(Pro-Hyp-Gly)_4_-Pro-Tyr-NH_2_  H-(Pro-Hyp-Gly)_4_-Pro-Met-Gly-Trp-Met-Gly-Ala-Lys-Gly-Pro-Hyp-Gly-(Pro-Hyp-Gly)_4_-Pro-Tyr-NH_2_  H-(Pro-Hyp-Gly)_4_-Pro-Nle-Gly-Trp-Nle-Gly-Ala-Hyp-Gly-Pro-Hyp-Gly-(Pro-Hyp-Gly)_4_-Pro-Tyr-NH_2_  Ac-(Gly-Pro-Hyp)_2_-Gly-Pro-Nle-Gly-Trp-Nle-Gly-Ala-Hyp-(Gly-Pro-Hyp)_2_-NH_2_ |

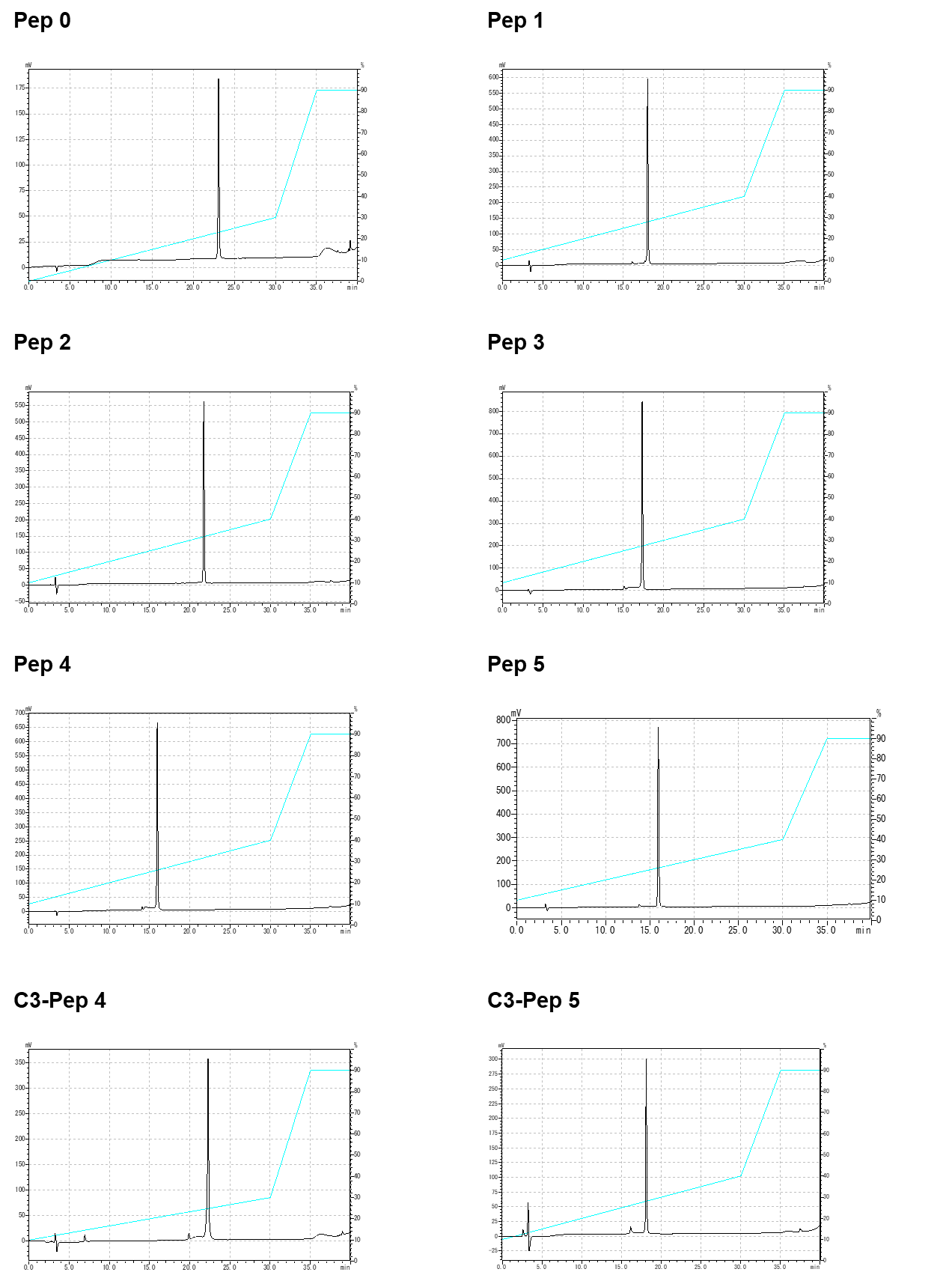

**Figure S2. Reverse-phase high-performance liquid chromatography (RP-HPLC) profiles of synthesized peptides.** Column: Cosmosil 5C18-ARII (4.6 i.d. × 250 mm). Gradient: 10%–40% CH_3_CN in 0.05% TFA/H_2_O. over 30 min at 60°C. Flow rate: 1.0 mL/min. Absorbance: 220 nm.

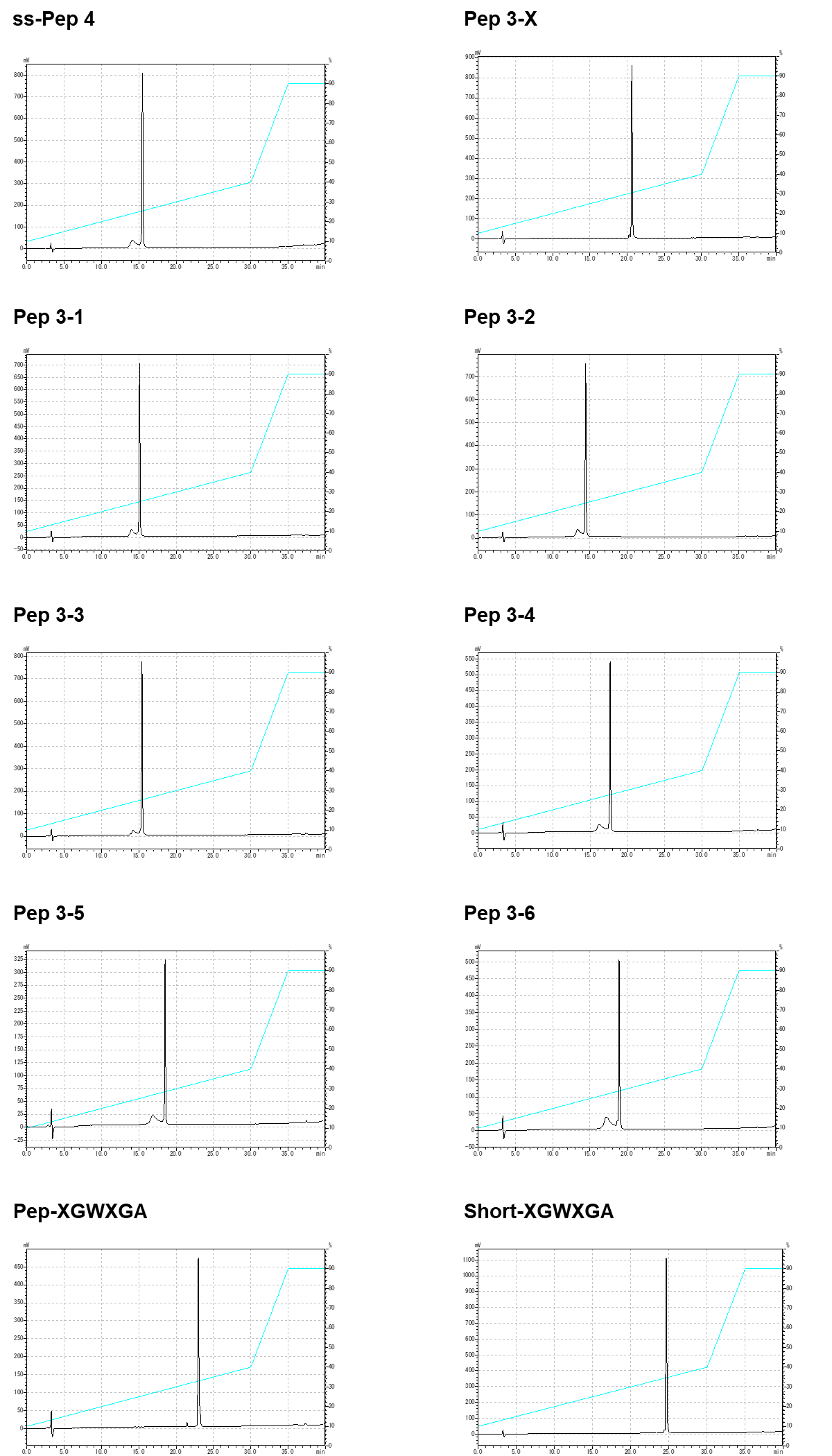

Figure S2. (*Continued*).

**Pep 0**

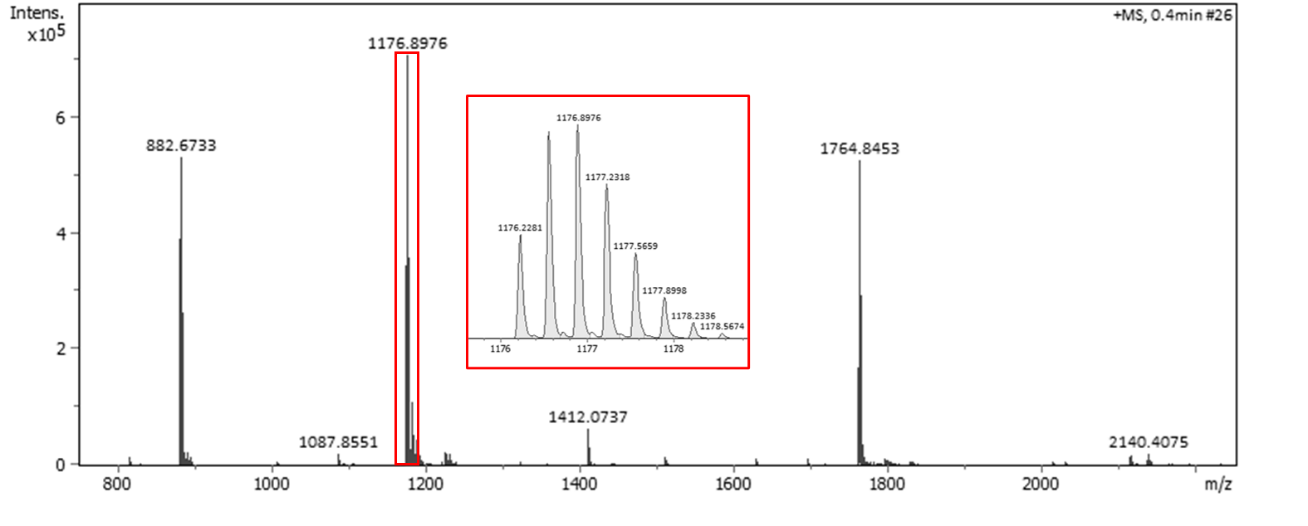

calcd. MS [M_m_ + 3H]^3+^: 1176.227	found: 1176.228

**Pep 1**

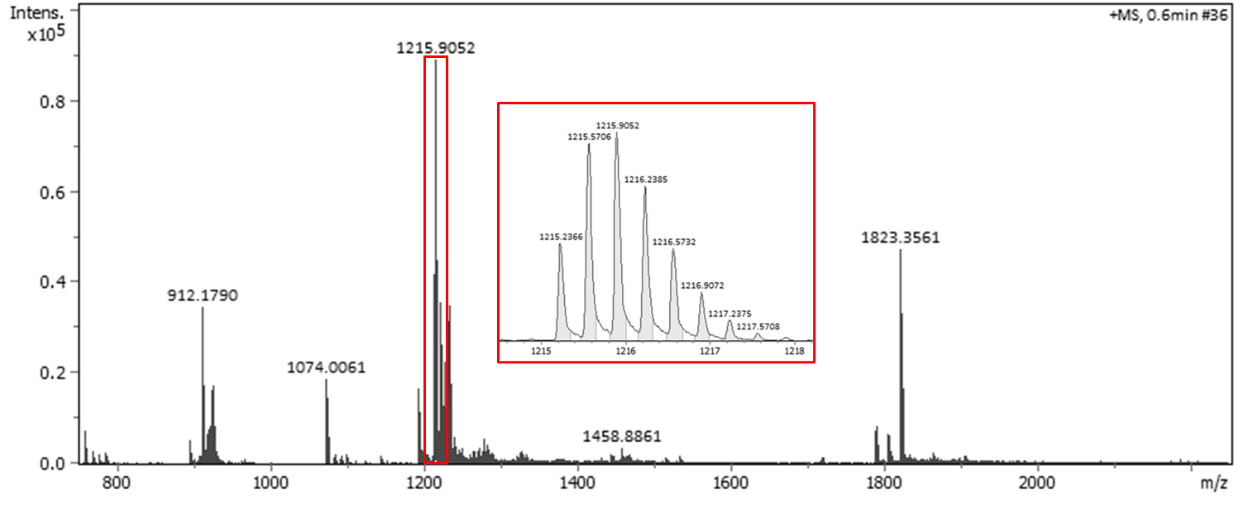

calcd. MS [M_m_ + 3H]^3+^: 1215.005	found: 1215.237

**Figure. S3. Mass spectrometry (MS) charts of the synthetic peptides.**

**Pep 2**

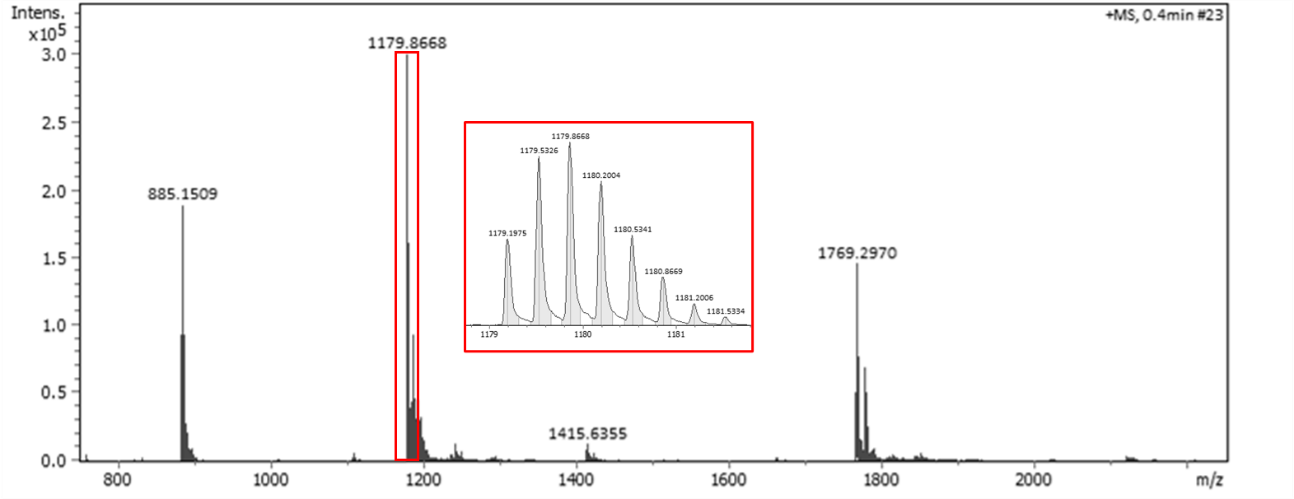

calcd. MS [M_m_ + 3H]^3+^: 1179.197	found: 1179.198

**Pep 3**

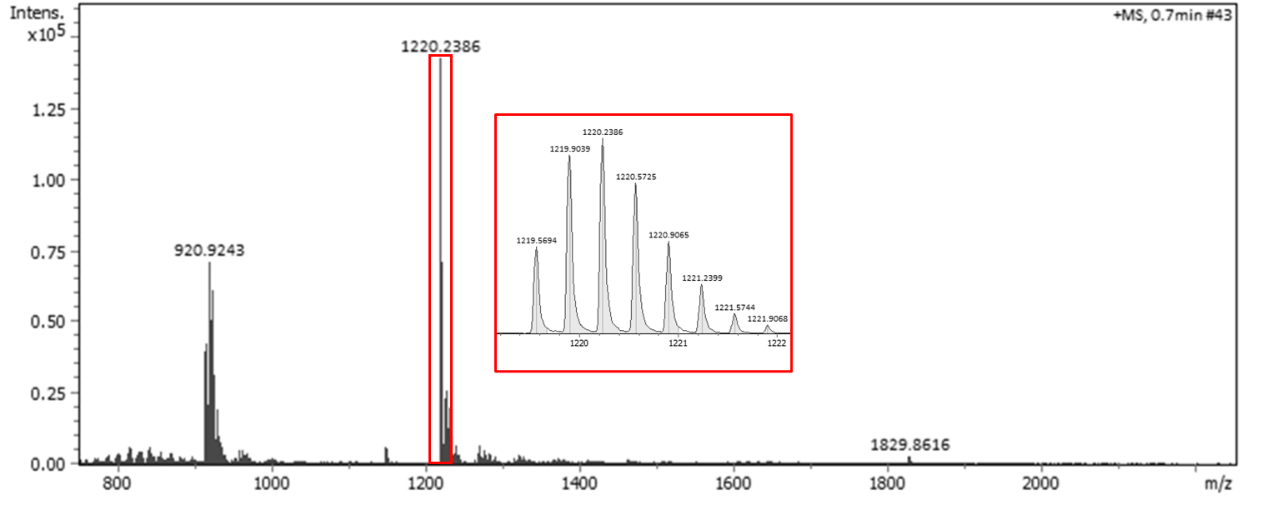

calcd. MS [M_m_ + 3H]^3+^: 1219.573	found: 1219.569

Figure S3. (*Continued*).

**Pep 4**

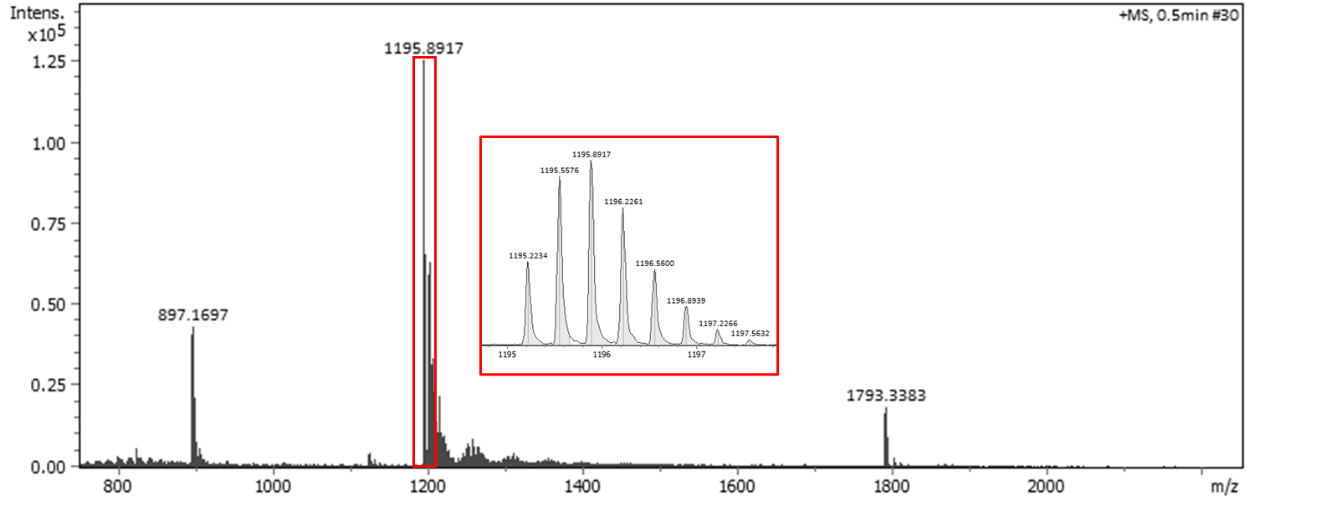

calcd. MS [M_m_ + 3H]^3+^: 1195.225	found: 1195.223

**Pep 5**

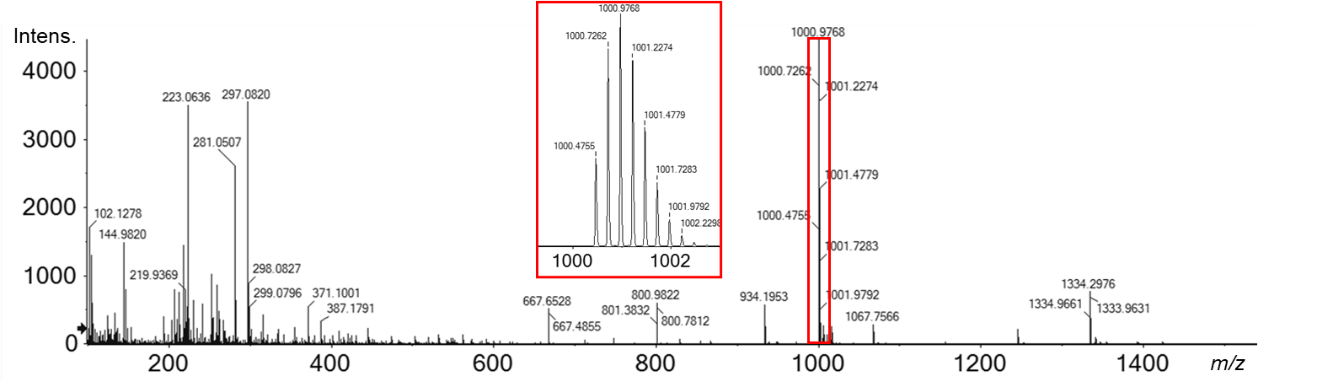

calcd. MS [M_m_ + 4H]^4+^: 1000.475	found: 1000.476

Figure S3. (*Continued*).

**C3-Pep 4**

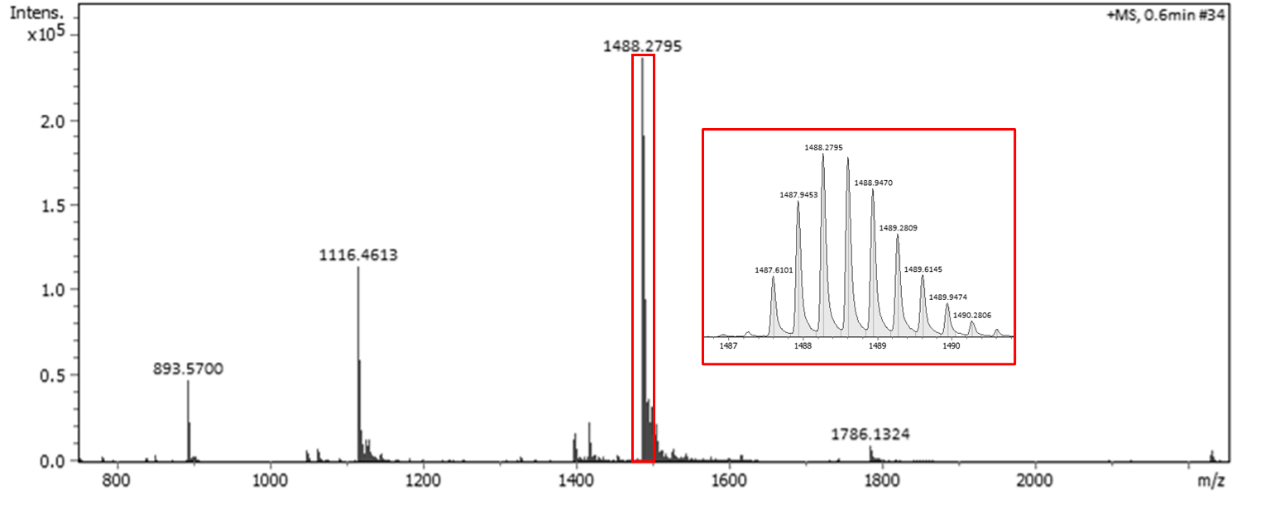

calcd. MS [M_m_ + 3H]^3+^: 1487.615	found: 1487.610

**C3-Pep 5**

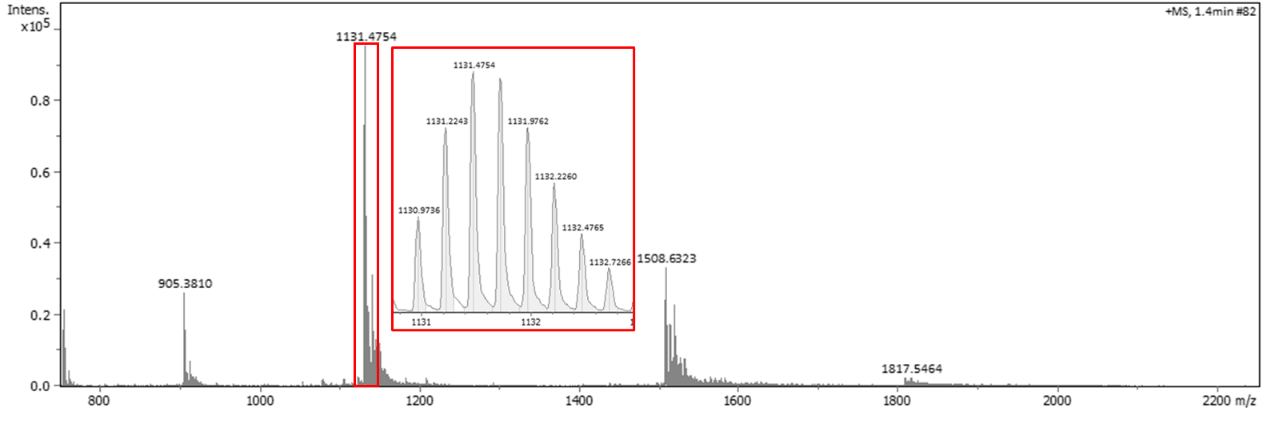

calcd. MS [M_m_ + 4H]^4+^: 1130.972	found: 1130.974

Figure S3. (*Continued*).

**ss-Pep 4**

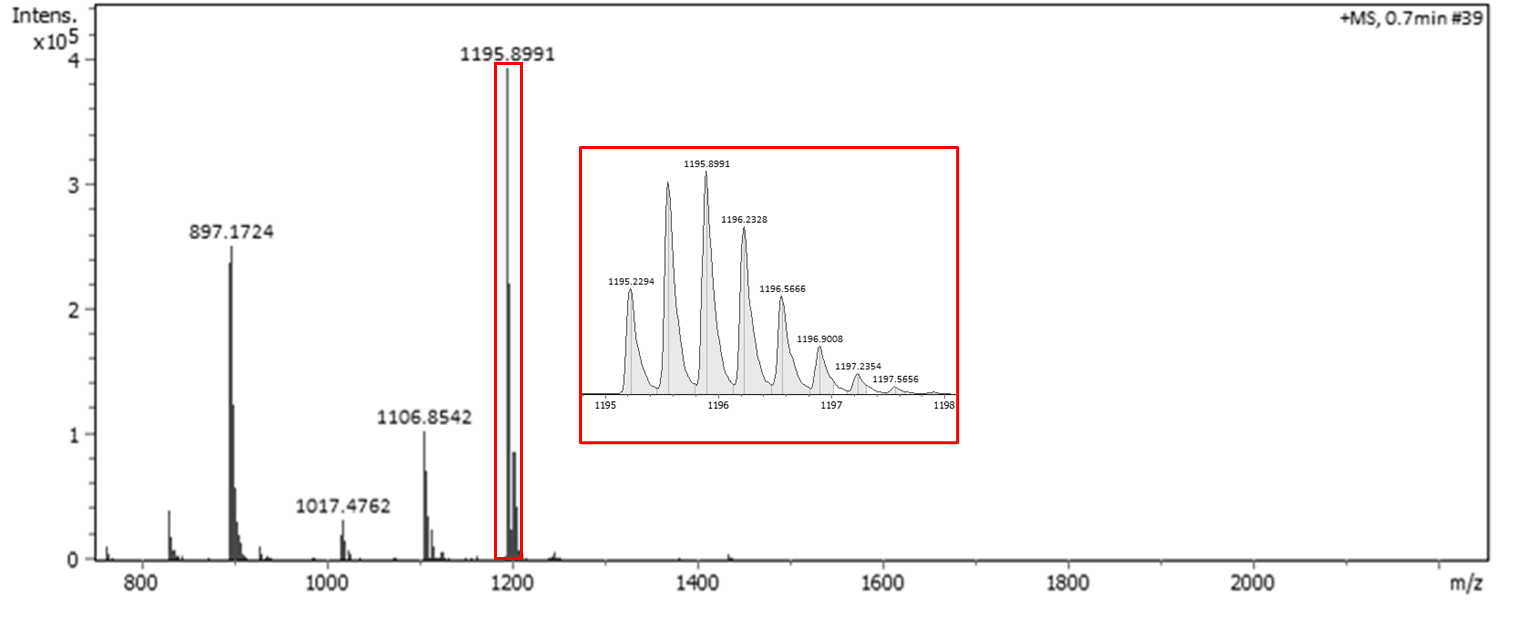

calcd. MS [M_m_ + 3H]^3+^: 1195.225	found: 1195.229

**Pep 3-X**

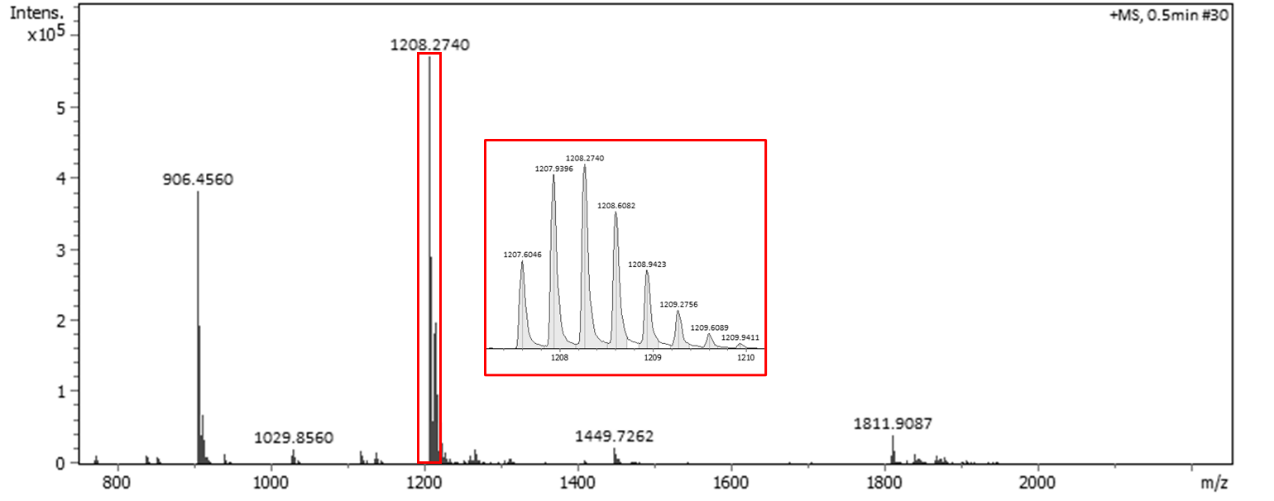

calcd. MS [M_m_ + 3H]^3+^: 1207.602	found: 1207.605

Figure S3. (*Continued*).

**Pep 3-1**

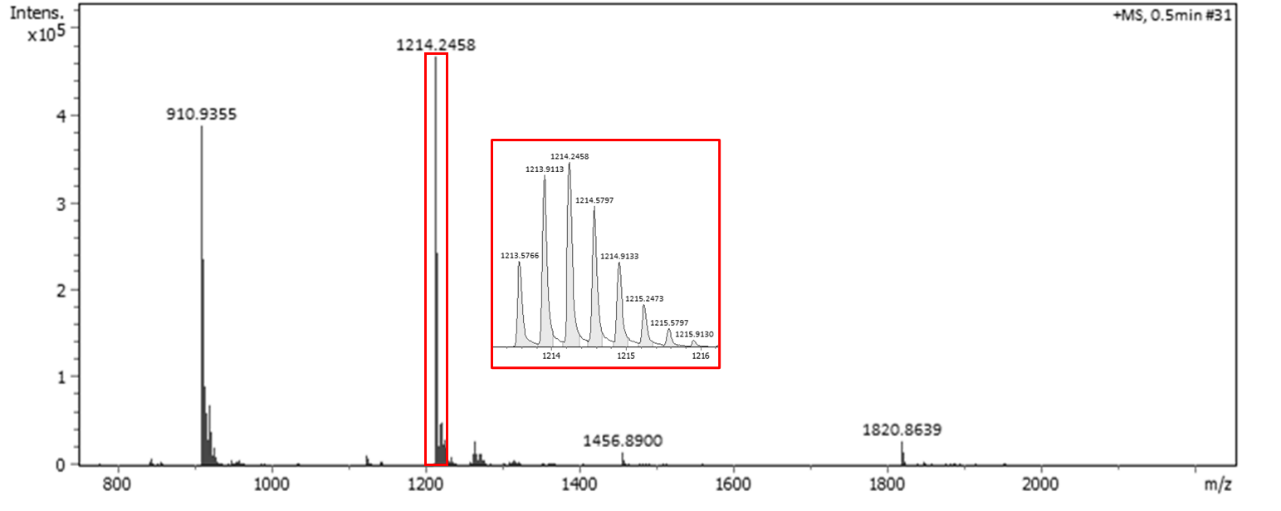

calcd. MS [M_m_ + 3H]^3+^: 1213.576	found: 1213.577

**Pep 3-2**

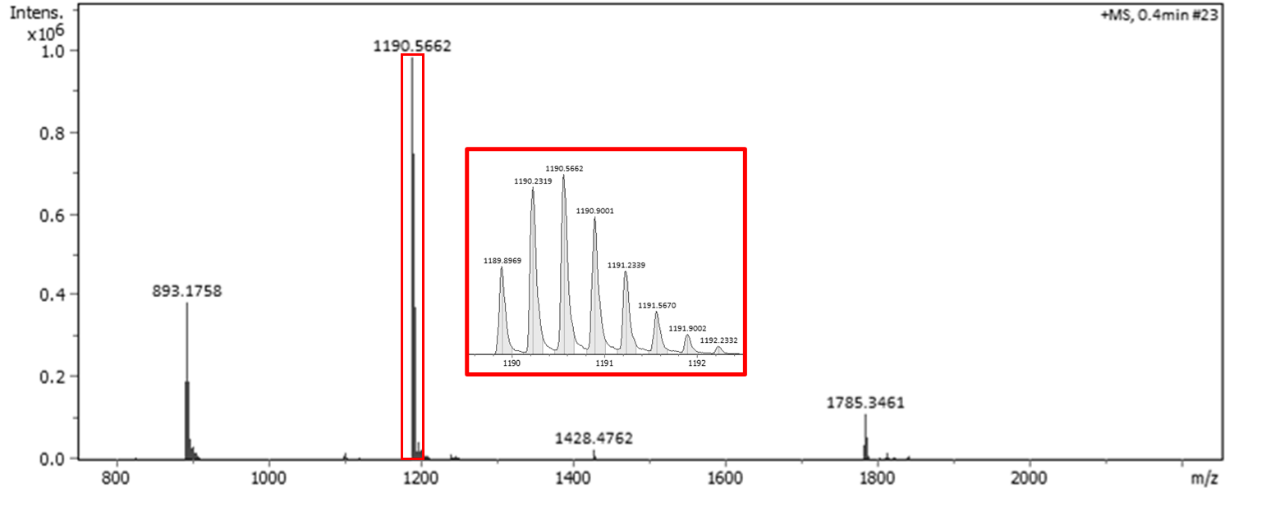

calcd. MS [M_m_ + 3H]^3+^: 1189.898	found: 1189.897

Figure S3. (*Continued*).

**Pep 3-3**

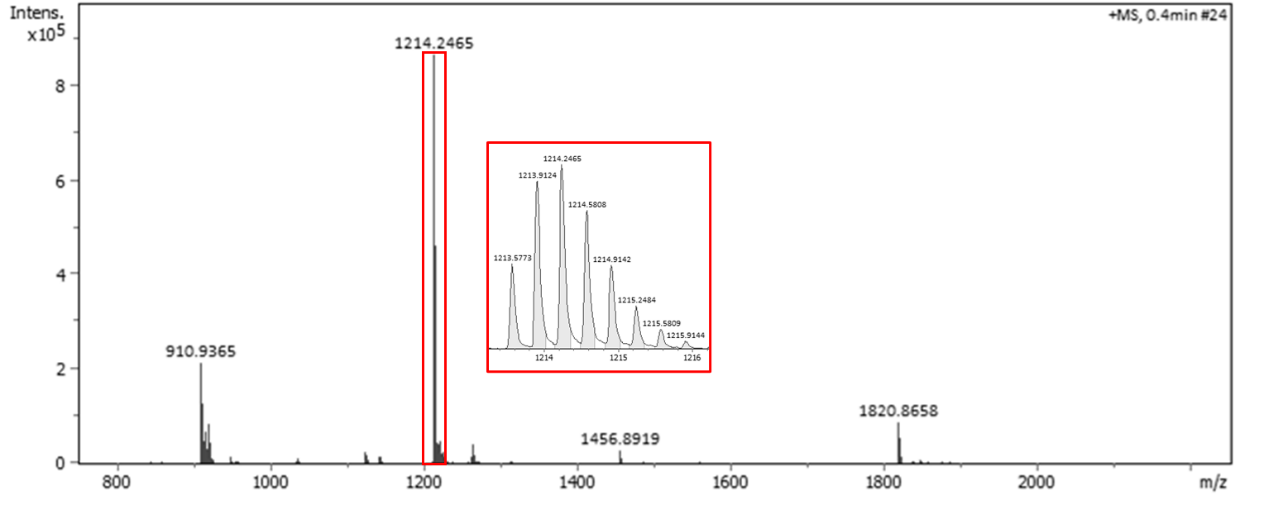

calcd. MS [M_m_ + 3H]^3+^: 1213.576	found: 1213.577

**Pep 3-4**

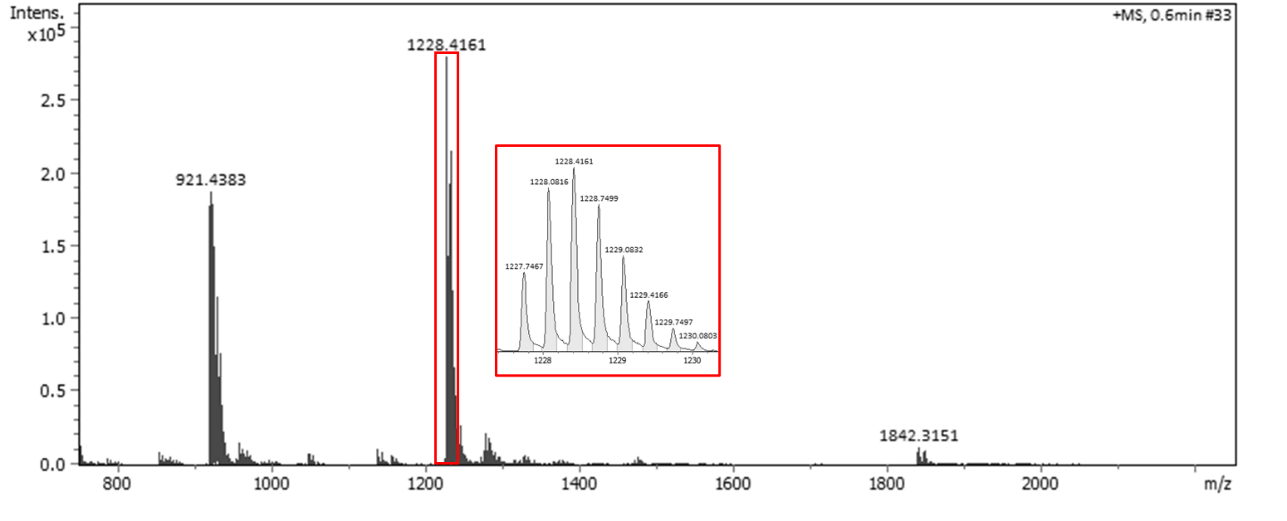

calcd. MS [M_m_ + 3H]^3+^: 1228.245	found: 1227.747

Figure S3. (*Continued*).

**Pep 3-5**

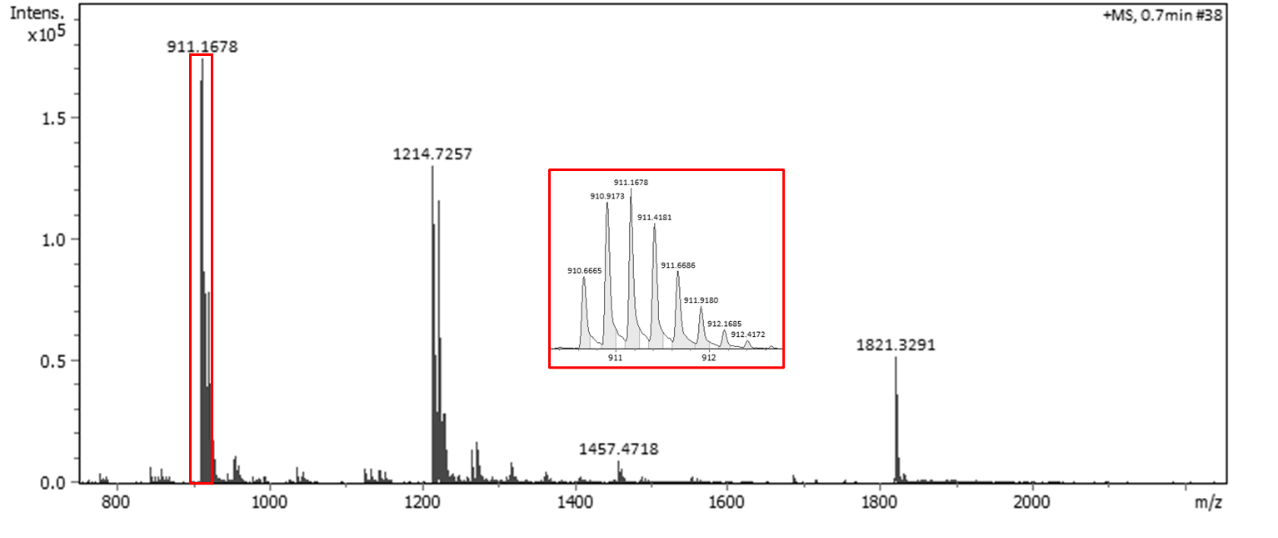

calcd. MS [M_m_ + 4H]^4+^: 911.170	found: 910.667

**Pep 3-6**

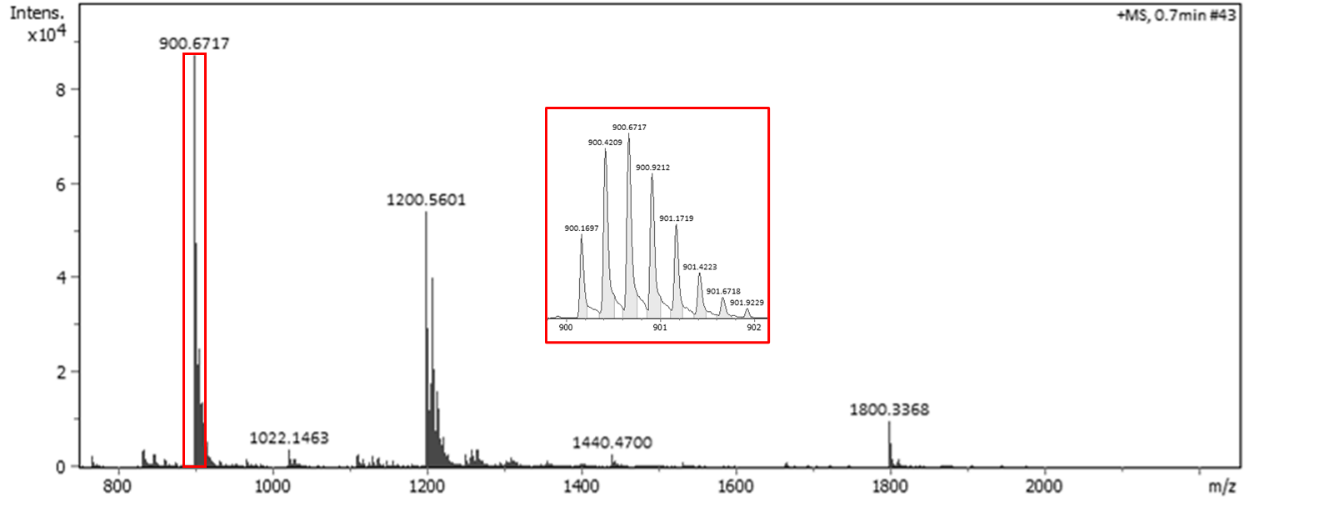

calcd. MS [M_m_ + 4H]^4+^: 900.170	found: 900.170

Figure S3. (*Continued*).

**Pep-XGWXGA**

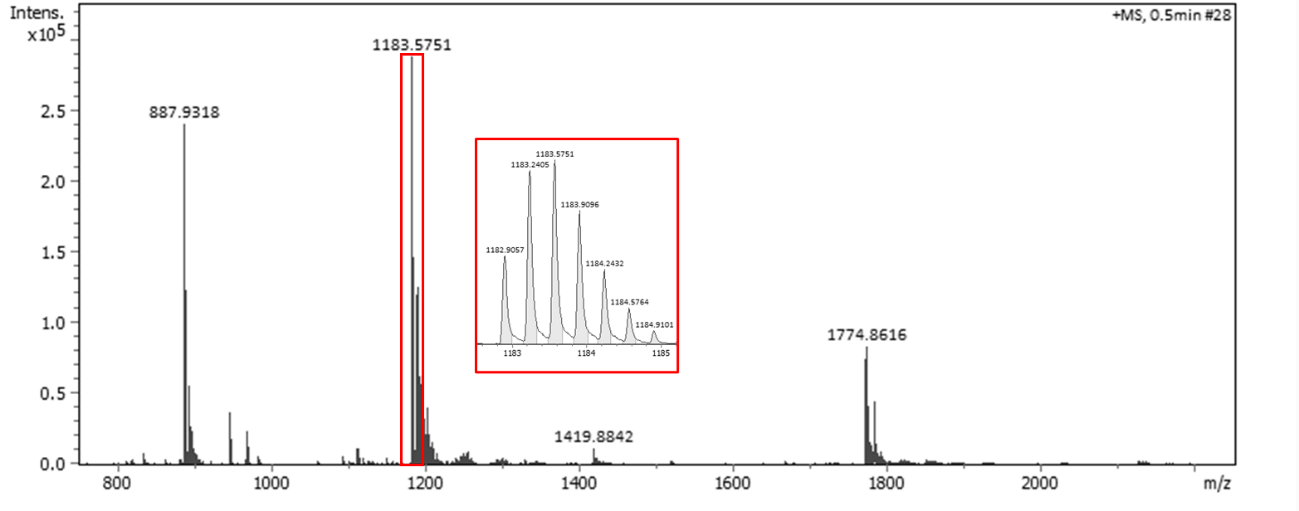

calcd. MS [M_m_ + 3H]^3+^: 1182.904	found: 1182.906

**Short-XGWXGA**

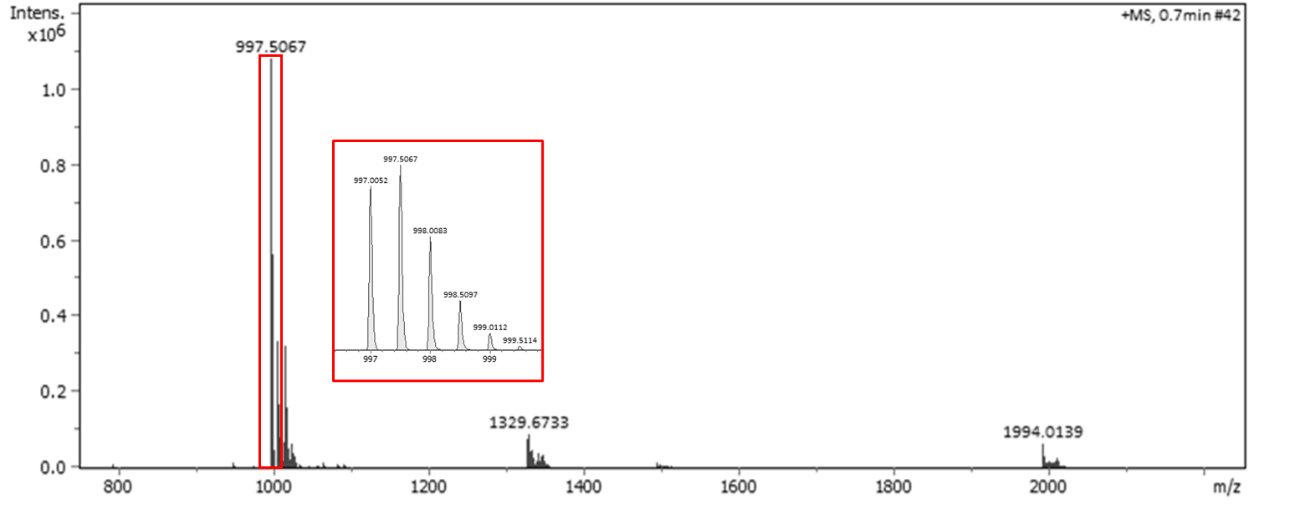

calcd. MS [M_m_ + 2H]^2+^: 996.995	found: 997.005

Figure S3. (*Continued*).

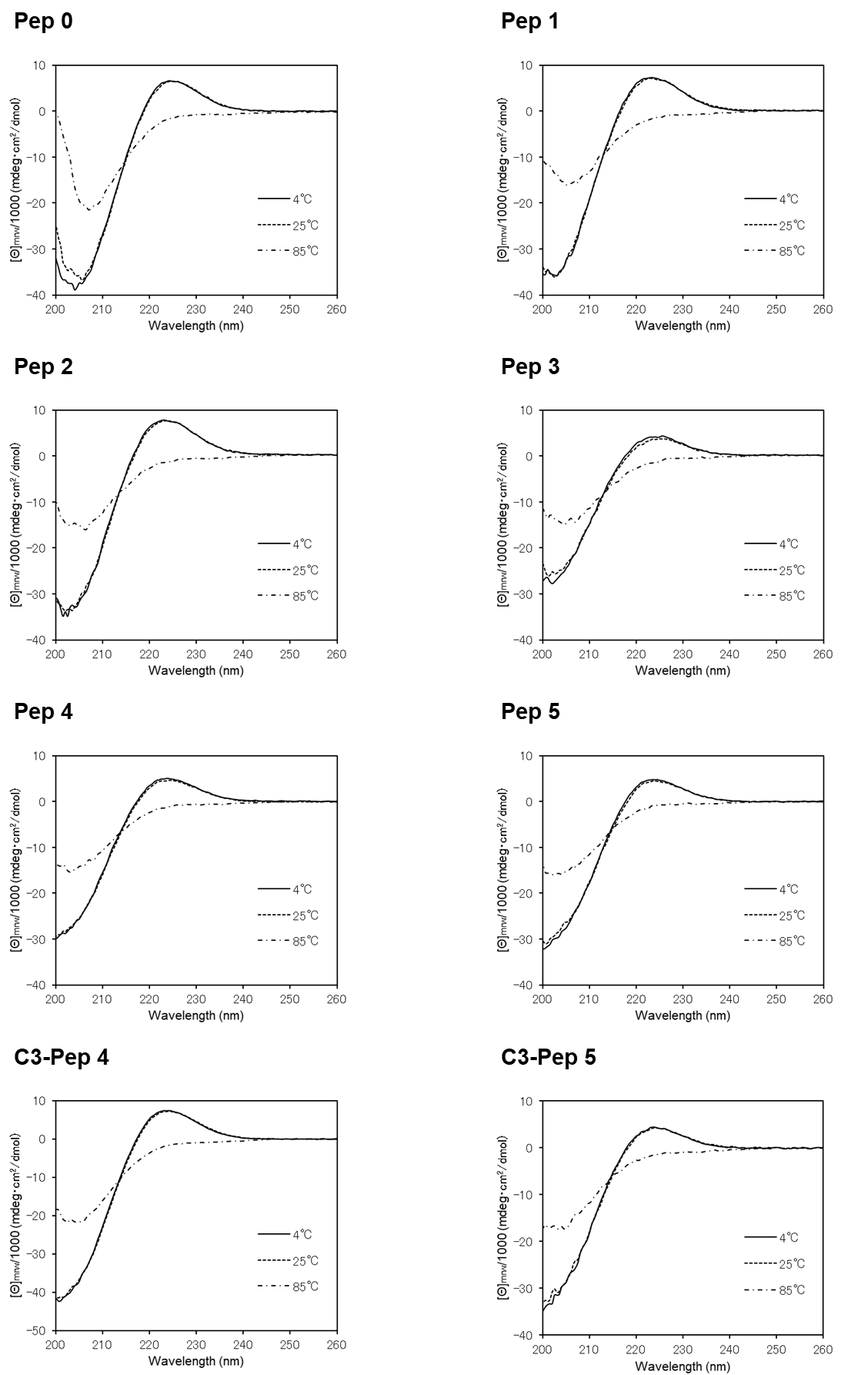

**Figure S4. Circular dichroism (CD) profiles of peptides at 4°C (solid line), 25°C (dashed line), and 85°C (dash-dotted line).** A positive maximum at 225 nm indicates a polyproline II-like helical structure (4).

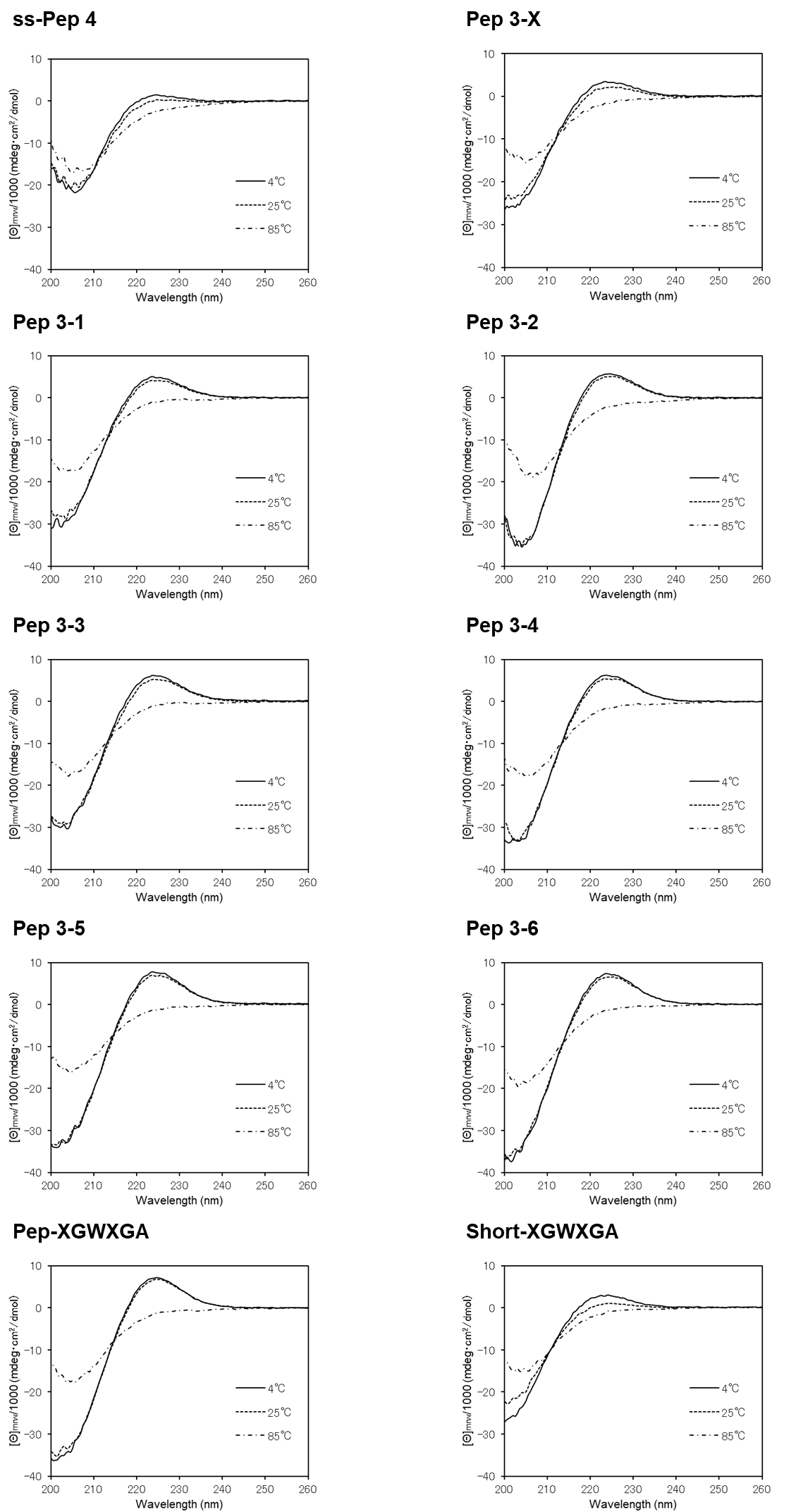

Figure S4. (*Continued*).

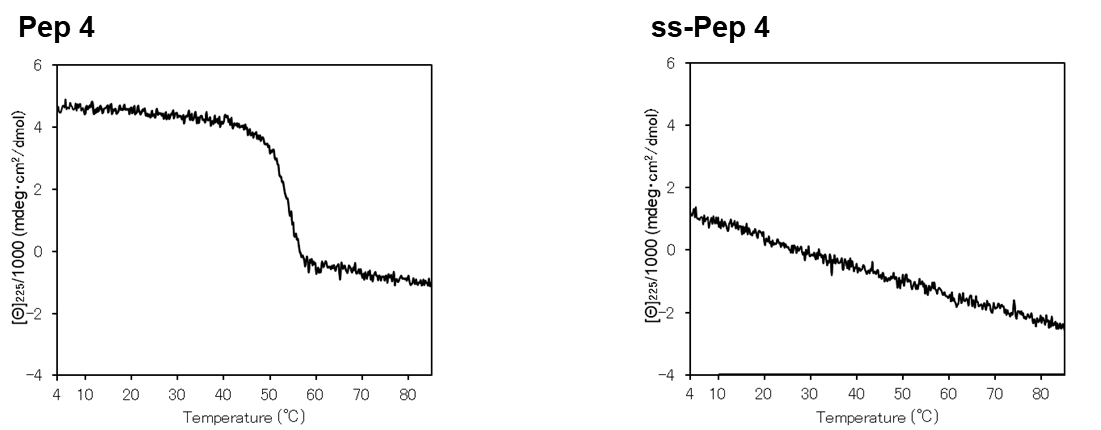

**Figure S5. CD spectra recorded at 225 nm from 4 to 85°C.** A cooperative decrease in the signal with increasing temperature indicates triple-helix melting, whereas a linear decrease reflects a single-chain polyproline II helix.

**Table S2. Predicted melting temperatures (*T*_m_) of triple-helical peptides containing Trp at Xaa3.**

| **Name** | **Sequence** | ***T*_m_ (°C) *^a^*** |
| --- | --- | --- |
| Clone 2    Clone 7    Clone 8 | H-(Pro-Hyp-Gly)_4_-Pro-Met-Gly-Trp-Trp-Gly-Pro-Ser-Gly-Thr-Hyp-Gly-(Pro-Hyp-Gly)_4_-Pro-Tyr-NH_2_  H-(Pro-Hyp-Gly)_4_-Pro-Met-Gly-Trp-Met-Gly-Ala-Lys-Gly-Arg-Hyp-Gly-(Pro-Hyp-Gly)_4_-Pro-Tyr-NH_2_  H-(Pro-Hyp-Gly)_4_-Pro-Met-Gly-Trp-Trp-Gly-Ala-Thr-Gly-Thr-Hyp-Gly-(Pro-Hyp-Gly)_4_-Pro-Tyr-NH_2_ | 9.7  32.9  17.2 |

*^a^* *T*_m_ value was predicted SCEPTTr (5).

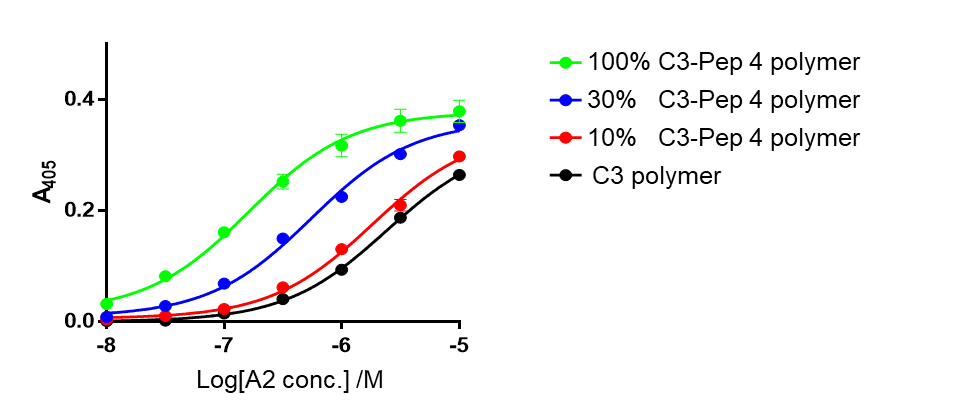

**Figure S6. Collagen VII A2 binding increased with the proportion of the C3-Pep 4 in the peptide polymer.** GST-collagen VII A2 was incubated with ELISA wells coated with peptide polymers (0.5 μg/well) at 25°C. Percentages of polymers represent ratios of the C3-Pep 4 weight to the total peptide weight (C3-Pep 4 and C3). Bound protein was detected using HRP-conjugated anti-GST antibody and ABTS substrate, and absorbance was measured at 405 nm (n = 3, mean ± S.D.).

**Table S3. Data collection and refinement statistics.**

|  | Human collagen VII A2 domain–Short-XGWXGA peptide complex |
| --- | --- |
| **Data collection**  Beamline  Detector  Wavelength (Å)  Space group  Unit-cell parameters  *a, b, c* (Å)  α, β, γ (º)  Unique reflections  Resolution (Å)  *R_meas_* (%)  CC_1/2_ (%)  Completeness (%)  Average *I/σ* (I)  Redundancy  **Refinement**  Resolution (Å)  Reflections  No. Atoms  Protein  Waters  SO4  *R_work_*  *R_free_*  R.M.S.D. from ideal  Bonds (Å)  Angles (º)  *B*-factors (Å^2^)  Ramachandran plot analysis (%)  Most favored  Allowed | SPring-8 BL45XU  EIGER X 16M  1.0  P2_1_  72.01, 37.17, 80.49,  90.00, 115.77, 90.00  65,405 (3,152)  40.39-1.47 (1.50-1.47)  30.6 (124.5)  99.3 (60.0)  99.3 (97.3)  15.6 (2.3)  6.8 (7.0)  40.39-1.47 (1.49-1.47)  65,389 (2,093)  3,745  606  10 (2 molecules)  0.176 (0.316)  0.217 (0.370)  0.008  1.034  19.00  96.17  3.83 |

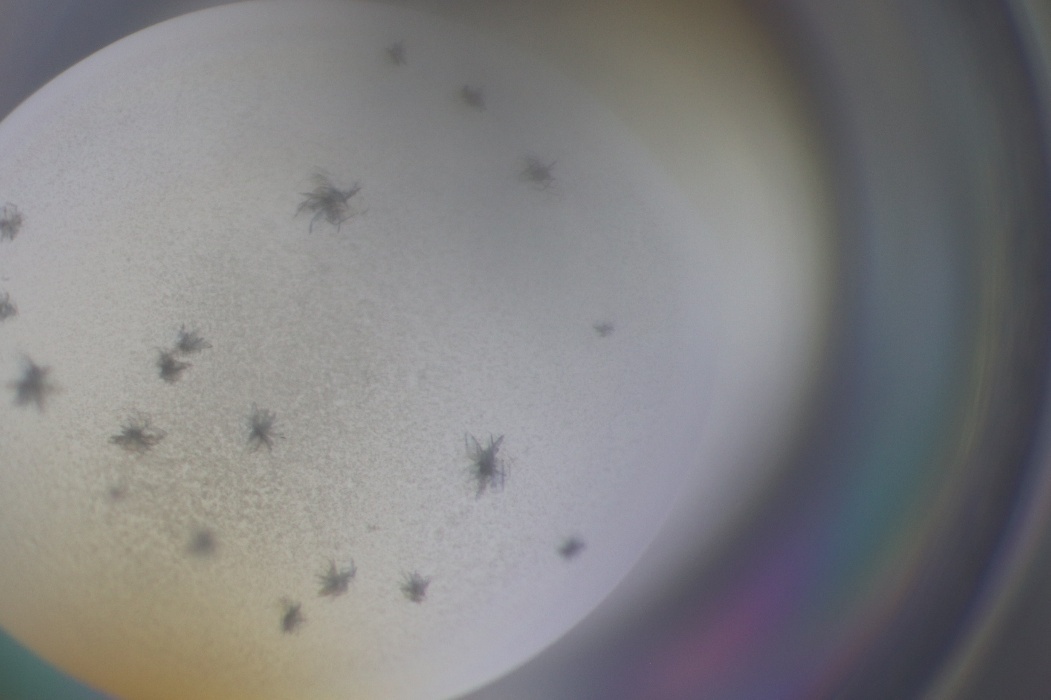

**Figure S7. Co-crystal of collagen VII A2 and Short-XGWXGA.** Purified A2 domain (243 μM) and Short-XGWXGA peptide (291 μM) were mixed, and crystallization screening was performed at 4°C. Diffraction-quality plate-like crystals were obtained in reservoir solution containing 26% (w/v) PEG 4000, 100 mM sodium acetate, and 100 mM ammonium sulfate at pH 4.8.

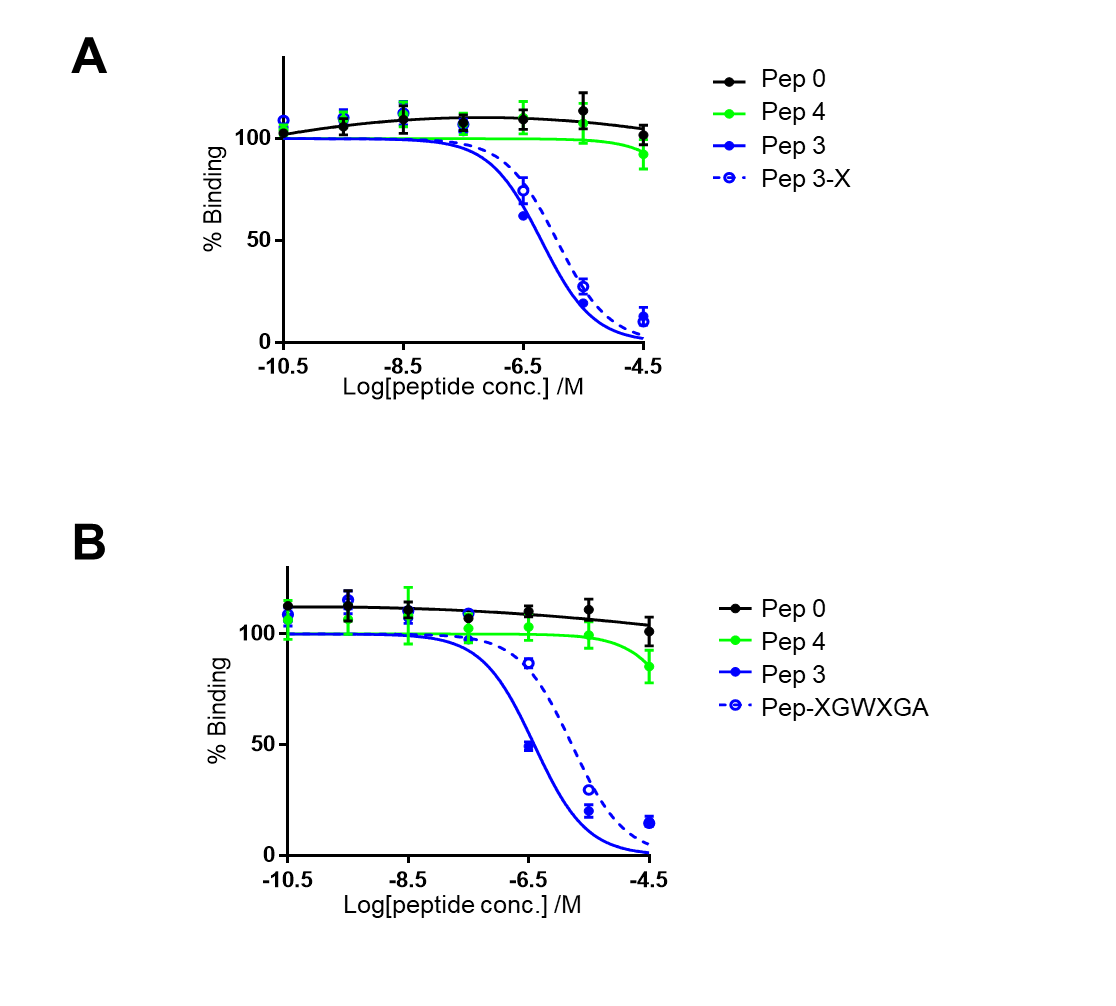

**Figure S8. Collagen VII A2 domain recognizes Nle-containing triple-helical peptide.** *A*, Binding of 10 nM GST-collagen VII A2 to wells coated with 100% C3-Pep 4 polymer (0.5 μg/well) was inhibited by Pep 3 (solid line) or Pep 3-X (dashed line). The experiment was conducted at 25°C (n = 3, mean ± S.D.). *B*, Binding of 10 nM GST-collagen VII A2 to wells coated with 100% C3-Pep 4 polymer (0.5 μg/well) was inhibited by Pep 3 (solid line) or Pep-XGWXGA (dashed line). The experiment was conducted at 25°C (n = 3, mean ± S.D.).

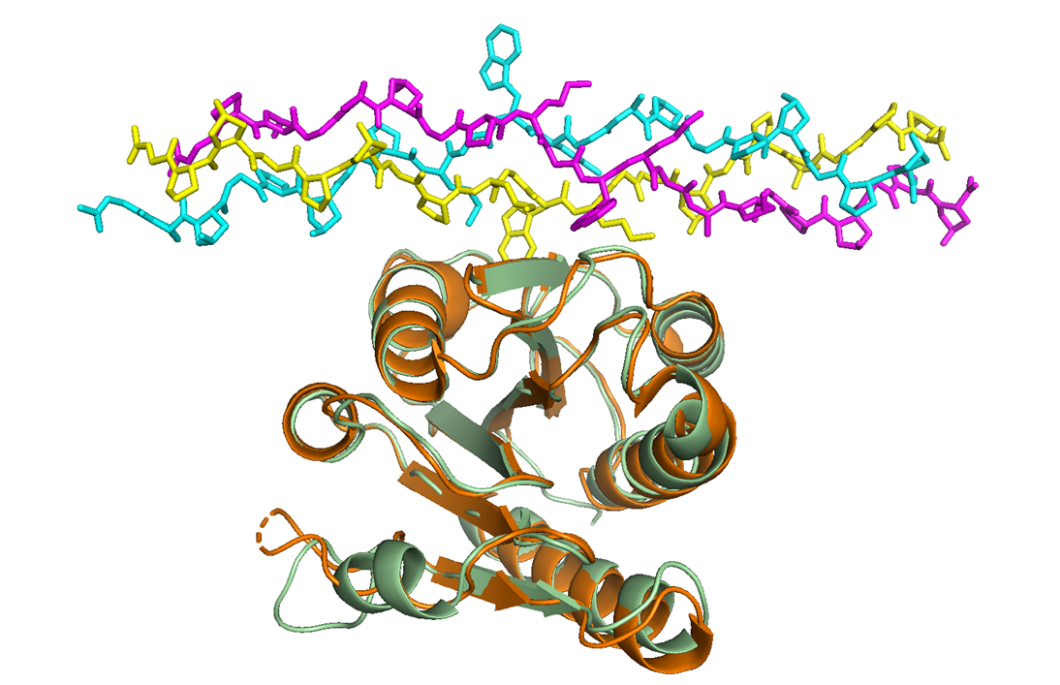

**Figure S9. Structural comparison of collagen VII A2 between the apo and complex forms.** Collagen VII A2 is shown as a cartoon model, and Short-XGWXGA as a stick model. The leading, middle, and trailing chains of Short-XGWXGA are colored cyan, yellow, and magenta, respectively. The A2 apo form (PDB: 6S4C) (6) is shown in orange, and the A2 complex form is shown in green. The RMSD is 0.506 Å.

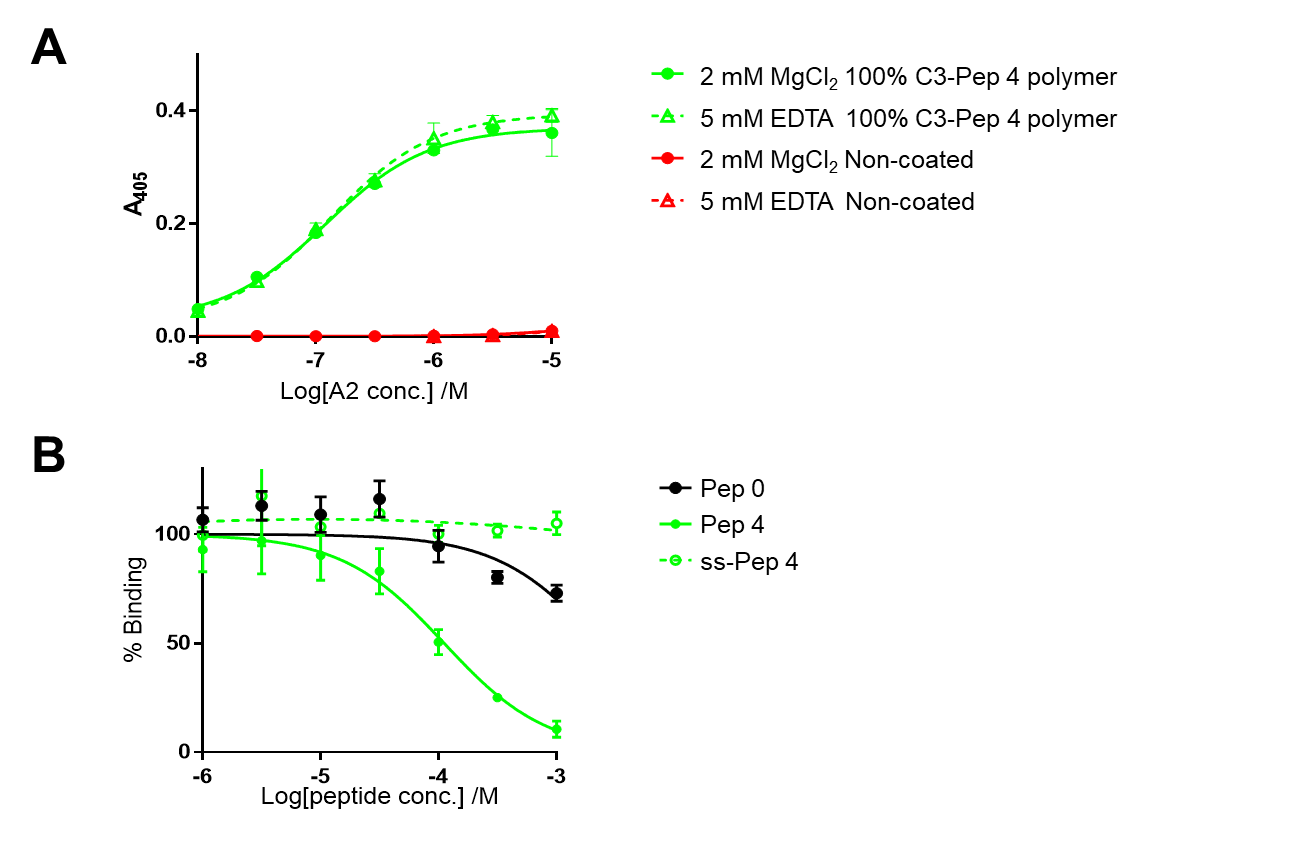

**Figure S10. Collagen VII A2 recognizes the collagen *triple-helical* peptides in a metal ion-independent manner.** *A*, GST-collagen VII A2 was incubated with ELISA wells coated with 100% C3-Pep 4 polymer (0.5 μg/well) at 25°C in the presence of MgCl₂ (solid line) or EDTA (dashed line). Bound protein was detected using HRP-conjugated anti-GST antibody and ABTS substrate. Absorbance was measured at 405 nm (n = 3, mean ± S.D.). *B*, Binding of 5 nM GST-collagen VII A2 to wells coated with 100% C3-Pep 4 polymer (0.5 μg/well) was inhibited by triple-helical Pep 4 (solid line) or single-stranded ss-Pep 4 (dashed line). Pep 4 that assumes triple-helical conformation at the assay temperature of 25°C, a stereoisomeric peptide containing D-Pro in place of L-Pro (ss-Pep 4), which fails to adopt a triple-helical conformation, was included as a competitor (Table S1). Circular dichroism analysis confirmed that ss-Pep 4 fails to form a triple-helical structure (Figure S4, Figure S5). Triple-helical Pep 0 served as a control. The experiment was conducted at 25°C (n = 3, mean ± S.D.).

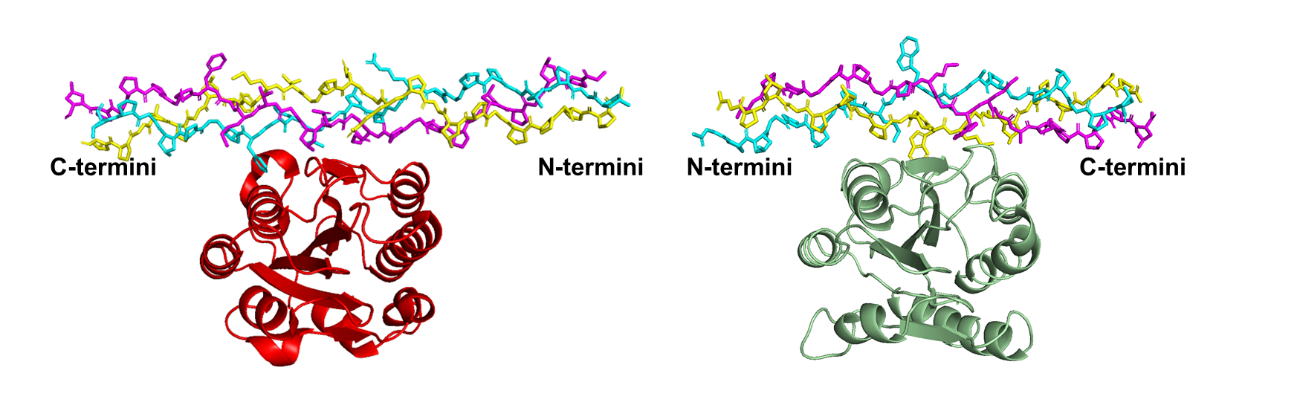

**Figure S11. Comparison of collagen-binding modes of collagen VII A2 and VWF A3.** The structure of VWF A3 (red) in complex with the triple-helical peptide was taken from a reported model (PDB: 4DMU) (7). The structure of collagen VII A2 (green) corresponds to the A2 domain extracted from the co-crystal structure determined in this study.

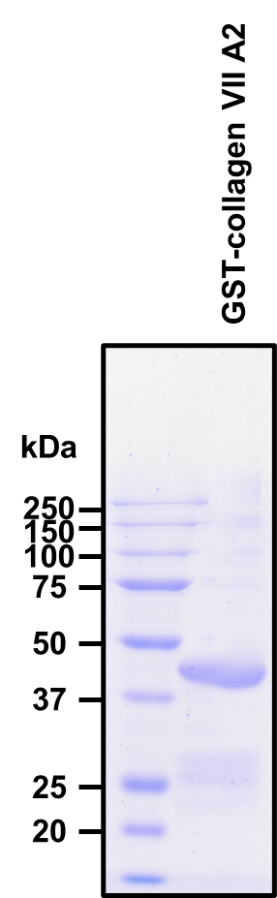

**Figure S12. SDS-PAGE analysis of purified GST-collagen VII A2 domain.** Proteins were separated on a 12% acrylamide gel, and bands were visualized by CBB staining. The molecular size of GST-collagen VII A2 domain is 48 kDa.
